# A structural biology community assessment of AlphaFold 2 applications

**DOI:** 10.1101/2021.09.26.461876

**Authors:** Mehmet Akdel, Douglas E V Pires, Eduard Porta Pardo, Jürgen Jänes, Arthur O Zalevsky, Bálint Mészáros, Patrick Bryant, Lydia L. Good, Roman A Laskowski, Gabriele Pozzati, Aditi Shenoy, Wensi Zhu, Petras Kundrotas, Victoria Ruiz Serra, Carlos H M Rodrigues, Alistair S Dunham, David Burke, Neera Borkakoti, Sameer Velankar, Adam Frost, Kresten Lindorff-Larsen, Alfonso Valencia, Sergey Ovchinnikov, Janani Durairaj, David B Ascher, Janet M Thornton, Norman E Davey, Amelie Stein, Arne Elofsson, Tristan I Croll, Pedro Beltrao

## Abstract

Most proteins fold into 3D structures that determine how they function and orchestrate the biological processes of the cell. Recent developments in computational methods have led to protein structure predictions that have reached the accuracy of experimentally determined models. While this has been independently verified, the implementation of these methods across structural biology applications remains to be tested. Here, we evaluate the use of AlphaFold 2 (AF2) predictions in the study of characteristic structural elements; the impact of missense variants; function and ligand binding site predictions; modelling of interactions; and modelling of experimental structural data. For 11 proteomes, an average of 25% additional residues can be confidently modelled when compared to homology modelling, identifying structural features rarely seen in the PDB. AF2-based predictions of protein disorder and protein complexes surpass state-of-the-art tools and AF2 models can be used across diverse applications equally well compared to experimentally determined structures, when the confidence metrics are critically considered. In summary, we find that these advances are likely to have a transformative impact in structural biology and broader life science research.

## Introduction

Proteins are the key molecules of the cell that are involved in all cellular processes. The three-dimensional (3D) shape of a protein provides critical information that can, among many things, be used to study protein interactions, functions and the impact of missense variation. While tremendous progress has been made in experimental approaches to determine protein structures, the experimentally determined structures of ~100,000 proteins (Burley et al., 2021) represent a very small fraction of the size and diversity of the universe of proteins. Protein structure prediction has been a fundamental challenge in bioinformatics for decades because accurate predictions could accelerate our understanding of protein structure/function relationships with vast impacts on the study of life. Since the first blind assessment of prediction methods much progress has been made (AlQuraishi, 2021), but none so radical as the one presented by Deepmind at CASP14 (Jumper et al., 2021). AlphaFold 2 (AF2) was shown to be able to predict the structure of protein domains at an accuracy matching experimental methods. Both the method and a database of 365,198 protein models have been released (Tunyasuvunakool et al., 2021) enabling the scientific community to better understand the accomplishments, abilities and limitations of AF2.

The accuracy of AF2 has been independently evaluated in blind assessments. Yet, many questions remain regarding the extent to which these approaches extend our coverage of structural biology, and the limitations of the AF2 method or structures derived from AF2 for applications in biology. Regarding coverage, previous attempts to generate “proteome-wide” structural models include those based on homology models such as the SwissModel Repository (SMR) (Bienert et al., 2017) and more recently, the modelling of known protein domains in the Pfam database (Mistry et al., 2021) using trRosetta (Yang et al., 2020). These represent prior benchmarks of large target coverage that can be useful to compare AF2 against.

In regards to the application of AF2 structures, it is noteworthy that it provides metrics of uncertainty (Jumper et al., 2021) that have been shown to reflect confidence in the structural assignment — potentially linked to protein disorder — and uncertainty for pair-wise residue distances. It is therefore important to assess whether AF2 structures and confidence metrics can be successfully integrated into and applied to critical structural biology tasks, such as functional classification, variant effects, binding site manipulation and modelling into new experimental data. (e.g. cryoEM). In addition to the prediction of individual protein structures, it has been recently shown that contact predictions can be used to simultaneously fold and dock proteins (Pozzati et al.) and early reports by the community have indicated that AF2 can predict the structure of complexes, which it was not initially trained to handle.

Here, we provide an evaluation and practical examples of applications of AF2 predictions across a large number of diverse structural biology challenges.

## Results

### Added structural coverage by AF2 predictions of model proteomes

The AF2 database released predictions of the canonical protein isoforms for 21 model species covering nearly all residues in 365,198 proteins representing around twice the number of experimental structures and 6 times the number of unique proteins with information deposited in the PDB. It is important to assess the extent by which AF2 predictions extend the structural coverage beyond previous attempts to generate “proteome-wide” structural predictions. We first compared the structures of 11 model species included in both SMR and AF2 databases. On average, AF2 adds an additional coverage of 44% of residues, ranging from 24% for *S. aureus* to 61% for *P. falciparum* (Fig 1A, residues). However, AF2 predicts structural information for all protein residues although not all are predicted with high confidence. For residues not present in SMR, we observe an average of 49.4% predicted with confidence (the predicted local distance difference test score, pLLDT>0.7), ranging from 27% for *P.falciparum* to 77% for *E.coli* (Fig 1A, AF residue confidence). A more stringent cut-off (pLDDT > 90) identifies on average 25% of residues with very high confidence predictions. On average, around 25% of the residues of the proteomes are covered by AF2 with novel (not present in SRM) and confident (pLDDT>0.7) structural predictions.

**Figure 1.**
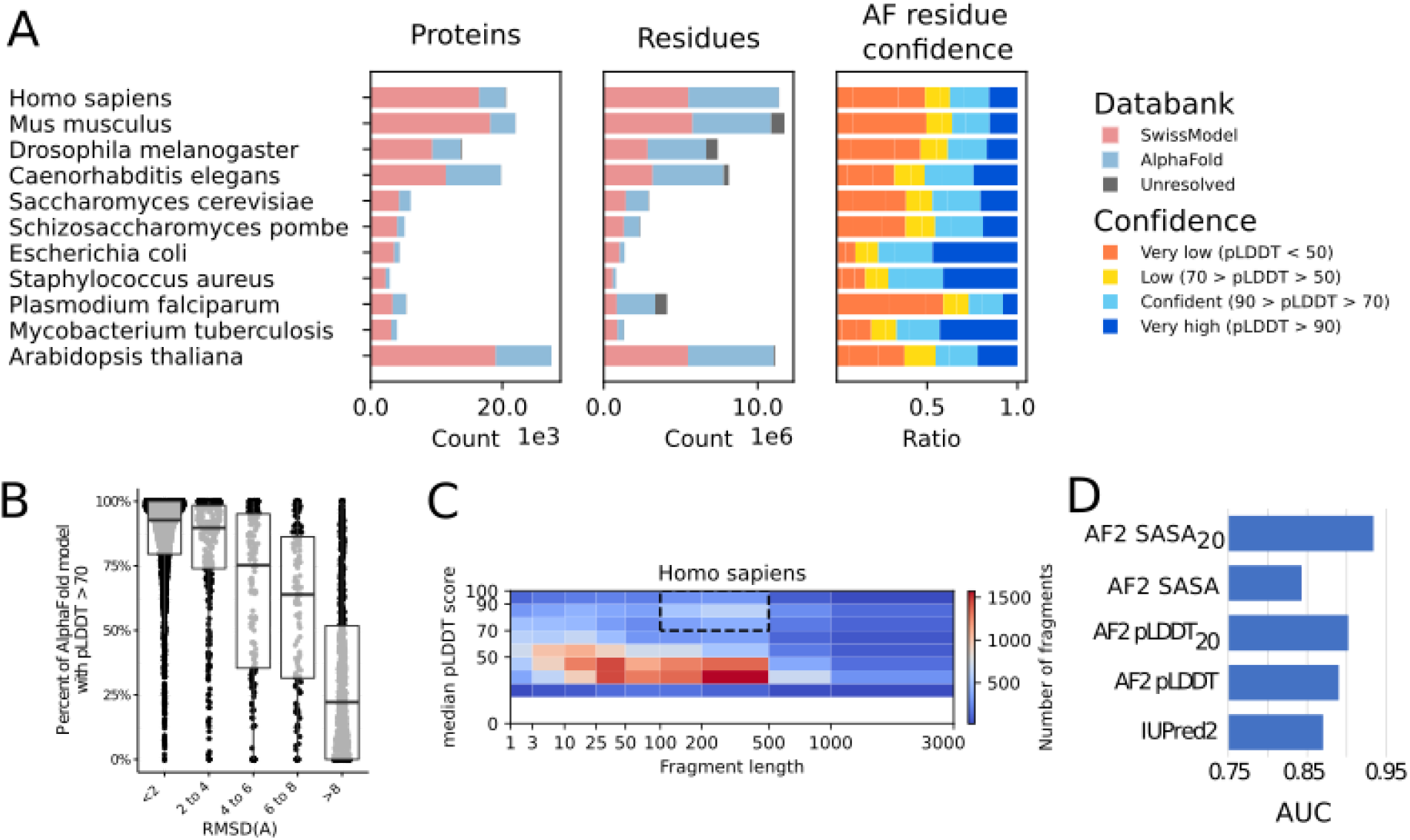
Additional coverage provided by AF2 predicted models. A) Added structural coverage (per-protein, left; per-residue, middle) and per-residue confidence of regions not covered by SMR (right) for 11 species shared between AF2 and SMR. B) Fraction of confident (pLDDT>70) residues per human AF2 model, binned by RMSD to the corresponding trRosetta-derived domain-level Pfam model. C) Median fragment length and median pLDDT score of human AF2-only regions. Highlighted area identifies high-confidence regions with domain-like length. D) Comparison of AF2 solvent accessible surface area (SASA; SASA_20_ adds 20-residue smoothing) and per-residue confidence (pLDDT; pLDDT_20_ adds 20-residue smoothing) against a state-of-the-art disorder prediction method (IUpred2).

We then compared AF2 predictions with those derived for Pfam protein domains (Mistry et al.,2021) using trRosetta (Yang et al., 2020). As there is only one trRosetta representative structure per domain family, we selected one species — human — and compared 3,035 AF2 models of 1,464 different Pfam domain families in human proteins with the representative trRosetta model. We find that these two approaches generally agree with around 50% of AF2 domain structures having RMSD<2Å to the generic trRosetta model (Fig S1A). We observe a correlation between the estimated accuracy of the AF2 model (pLDDT) and the RMSD from the trRosetta model (Fig 1B and Fig S1B-C). For example, the AF2 models with an RMSD below 2Å to the trRosetta model have, on average, more than 90% of their residues with a pLDDT above 70 (Fig 1B). We also examined the variability of domain structure for 273 domain families with 3 or more instances in the human proteome (Fig S2), observing that 70% of domain instances are within one standard deviation of the mean RMSD for their domain family. Together these results indicate that for at least 50% of human Pfam domains the trRosetta Pfam model was already likely to be accurate.

We assessed the confidence and length of AF2 contiguous regions not covered in SMR to identify regions that may correspond to novel structures of folded domains rather than short termini or interdomain linkers. The distribution of median confidence scores of a fragment vs. fragment length shows an enrichment for high-confidence predictions with 100–500 residue length (Fig 1C and Fig S3) consistent with the size of a typical protein domain (Wheelan et al.,2000). This relation can be observed for all species except *S. aureus* (Fig S3). We identified, across the 11 species, 18,429 contiguous regions that are “domain like” (100 ≤ len < 500) with confident predictions (pLDDT>70) that have no model in SMR. These are provided for human in Table S1.

While around half of the residues in AF2 predictions of the 11 model species are of low confidence, many of these may correspond to regions without a well-defined structure in isolation. About one-third of the human proteome corresponds to such intrinsically disordered proteins or protein regions (IDPs/IDRs) and it was shown that regions with low pLDDT are often IDRs (Tunyasuvunakool et al. 2021). We benchmarked AF2 against IUPred2 (Mészáros et al., 2018), a dedicated state-of-the-art disorder prediction method (Fig 1C), using a set of regions annotated for order/disorder (Table S2). In addition to using the per residue confidence scores (pLDDT), we also tested relative solvent accessible surface area (SASA) of each residue and smoothed versions of the pLLDT and SASA (Fig 1D and Fig S4). We observed that pLDDT and window averages of pLLDT or SASA clearly outperform a dedicated disorder predictor with a 20 residue smoothing window providing the best performance. This result strongly indicates that the AF2 low confidence predictions are significantly enriched for IDRs. To facilitate the study of human IDRs we provide these predictions for human proteins in supplementary results (Supplementary dataset 1) and we also integrated these measures into ProViz (Jehl et al.,2016), an online protein feature viewer available at http://slim.icr.ac.uk/projects/alphafold?page=alphafold_proviz_homepage.

In summary, these results indicate that prior to AF2 there was already significant structural coverage that could be derived by existing experimental and computational approaches. Nevertheless, AF2 provides substantial novel structural information to a model proteome, including high confidence structures and disorder predictions. These could be of potential future interests, including for the study of condensation of low complexity or intrinsically disordered proteins.

### Characterization of structural elements in AF2 predicted models across 21 proteomes

The AF database contains full proteome structure predictions across 21 species and is scheduled to grow to cover over 100 million proteins. The AF2 high confidence predictions are likely to contain a large number of rarely studied folds and structural elements that may not have been extensively seen in experimental structures. Due to the presence of low confidence regions in the AF proteins we first performed trimming to split each AF predictions into smaller high confidence units (see Methods). We then performed a global comparison of structural elements between the 365,198 proteins in the AF database and 104,323 proteins from the CASP12 dataset in the PDB. We applied the Geometricus algorithm (Durairaj et al., 2020) to obtain a description of protein structures as a collection of discrete and comparable shape-mers, analogous to k-mers in protein sequences and words in natural language processing (NLP). We then obtained a matrix of such shape-mer counts for all proteins. To identify substructures that may be novel or poorly represented, we clustered the proteins according to their shape-mer counts using Non-negative Matrix Factorization (NMF) (see Methods). The clustering identified 250 groups of proteins, dubbed “topics”, (Supplementary Dataset 2) with characteristic combinations of shape-mers. These characteristic shape-mers could include small structural elements such as repeats or the specific arrangements of ion binding sites, or larger structural elements that could define specific folds.

For visualisation we performed a t-SNE dimensionality reduction where proteins composed of similar shape-mers are expected to group closer together (Fig 2). In line with this, the shape-mer representation of AF2 proteins can predict the corresponding PDB protein entries with high accuracy (ROC-AUC of 0.95 using the cosine similarity of the shape-mer vector). Additionally, the 20 most common superfamilies, predicted from sequence, tend to be placed together in this low dimensional representation of shape-mer space. Importantly, even for superfamilies with some experimental structural characterization, AF2 predictions can greatly expand coverage such as for the transmembrane G-protein coupled receptors (in dark blue).

**Figure 2.**
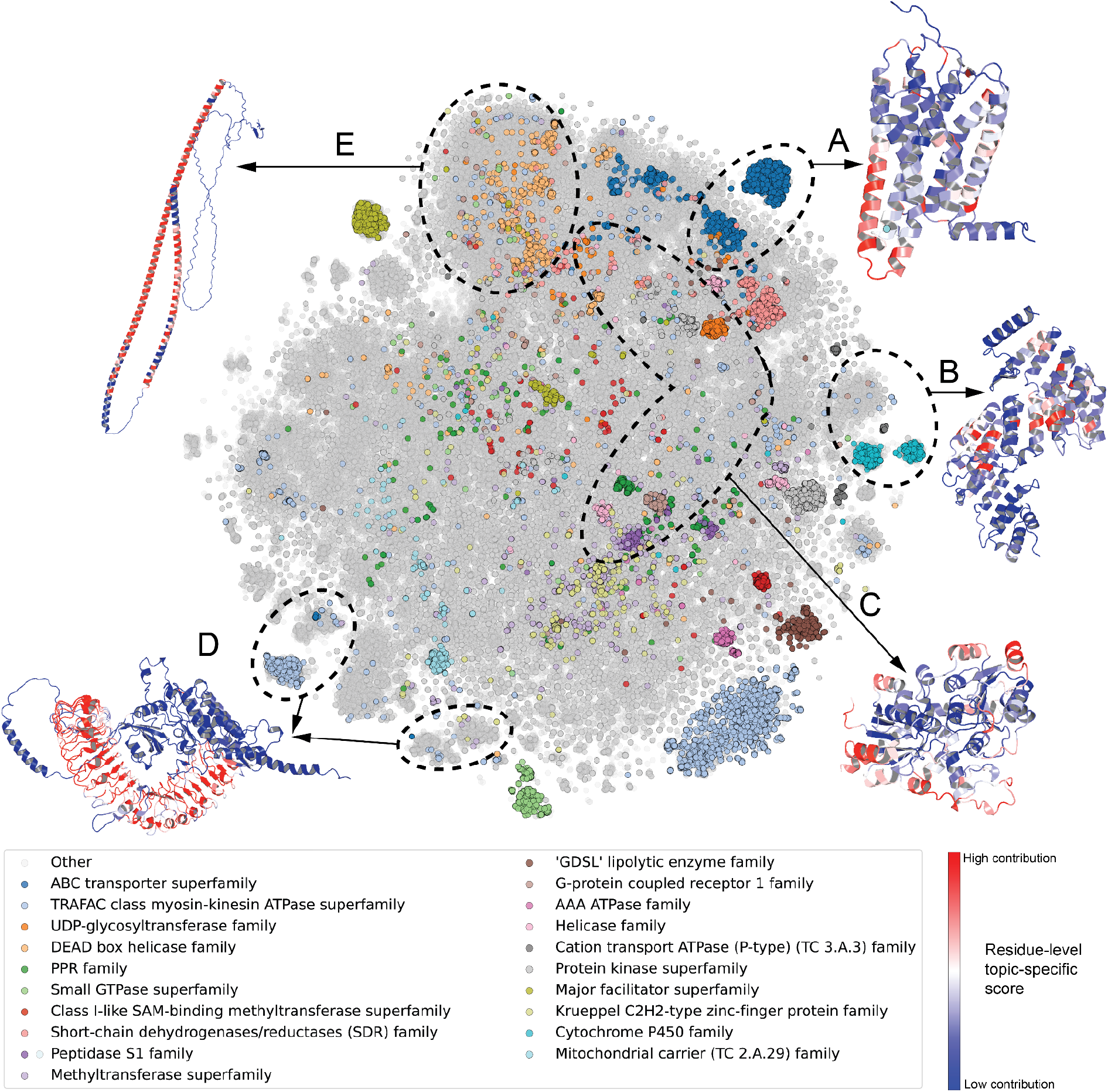
The space of characteristic structural elements in AlphaFold 2 structural models for 21 species — the 20 most common superfamilies are colored. Structures with similar structural elements are placed closer together in the figure. Five shape-mer groups (i.e. topics) discussed in the text, consisting of mainly (>90%) AlphaFold proteins as opposed to PDB proteins, are labeled A–E and a representative structure is depicted for each. Residues in the representative structures are colored according to their contribution to the topic under consideration — red residues have the highest contribution, while blue residues are specific to the example and not to the topic.

Out of 250 total topics, consisting of proteins with characteristic structural elements, we selected five examples that were almost exclusively (>90%) composed of structures derived from AF2. We illustrated these with a representative structure in Fig 2. Examples include 4,192 proteins annotated as G-protein coupled olfactory or odorant receptors (Pfam PF13853), 97% of which are mammalian (Fig 2A, topic 88, Fig S5A). Other examples include a group of primarily (94%) plant proteins annotated as PCMP-H and PCMP-E subfamilies of the pentatricopeptide repeat (PPR) superfamily (Fig 2B, Topic 60, Fig S5B); a group of heterogeneous structures mostly (>75%) annotated as ATP or ion binding (Fig 2C, Topic 150, Fig S5C); groups of proteins with leucine-rich repeats (Fig 2D, topic 16, Fig S5D); and long α-helical constructs (Fig 2E, Topic Helix, Fig S5E). For the PCMP-H and PCMP-E subfamilies (Fig 2B) there are no known experimental structures mapped. AF2 predictions could help elucidate the structural peculiarities of these subfamilies, including the mechanism of RNA recognition and binding for PCMP-H and PCMP-E proteins.

Finally, we also considered protein groups consisting primarily of PDB proteins to study why AF2 proteins are absent here. In some cases this seemed to be due to the limited number of species and proteins covered by the current AF2 database. For example, Topic 7 consists of 64% Enterobacteria phage T2 “Glycosyl hydrolase 24” family proteins, a species not yet modeled by AlphaFold. Topics 209 and 113 consist of immune response proteins such as immunoglobulins and T-cell receptors, mainly from the PDB. As many of these antibodies are under intense study, there are many more PDB structures (based on multiple individuals and antibody-drug research) than the actual number of such proteins in the respective UniProt proteomes. Topic 38 consists of short fragments of PDB structures, with an average length of 12 +/− 4 residues - there are no AF2 proteins here as AlphaFold models the entire structure instead of returning fragments.

Even for the 21 proteomes modelled with AF2 to date, this analysis already indicates the potential for a large expansion of structures for rarely studied folds which may lead to the identification of novel structural elements. These analyses indicate the areas of protein structure space that may be of highest interest for experimental characterization and should facilitate the study of the evolution of protein folds.

### Application of AF2 models for structure based variant effect prediction

A protein structure facilitates the generation of hypotheses regarding the impact of missense mutations or the rationalization of experimentally measured consequences. Conversely, an agreement between expected and observed impact of mutations provides confidence in the accuracy of a structural model. We obtained two independent compilations of experimentally measured impacts of protein mutations on protein function: 1) a compilation of measured changes in stability upon mutations (Nikam et al., 2021; Xavier et al., 2021); and 2) a compilation of deep mutational scanning (DMS) experiments (Dunham and Beltrao, 2021) that can measure the outcome of any possible single point mutation on a large proportion of all protein positions.

The deep mutational experimental measurements are available for 33 proteins with 117,135 total mutations. From these we could obtain experimentally derived models for 31 and AF2 models for all 33. We then used 3 structure-based variant effect predictors (FoldX (Delgado et al., 2019), Rosetta (Smith and Kortemme, 2011) and DynaMut2 (Rodrigues et al., 2021)) to compare the DMS measurements with predicted impacts using either AF2 or experimentally derived structures. While the correlation estimates between experimental and predicted impact of mutations varied across the proteins, those derived with the AF2 models were consistently matching or better than those derived from experimental models (Fig 3A, 3B and Fig S6). Regions with confidence scores lower than pLDDT<0.5 result in lower concordance (Fig 3A) but remarkably, restricting to protein regions without an experimental model can still lead to correlations that are comparable to those observed in experimental structures (Fig 3B). As low AF2 confidence scores are enriched for intrinsically disordered protein regions it is possible that the poor correlation in low confidence regions is in part due to higher tolerance to protein mutations. In line with this, we observed an average higher tolerance to mutations in low confidence regions (Fig 3C).

**Figure 3.**
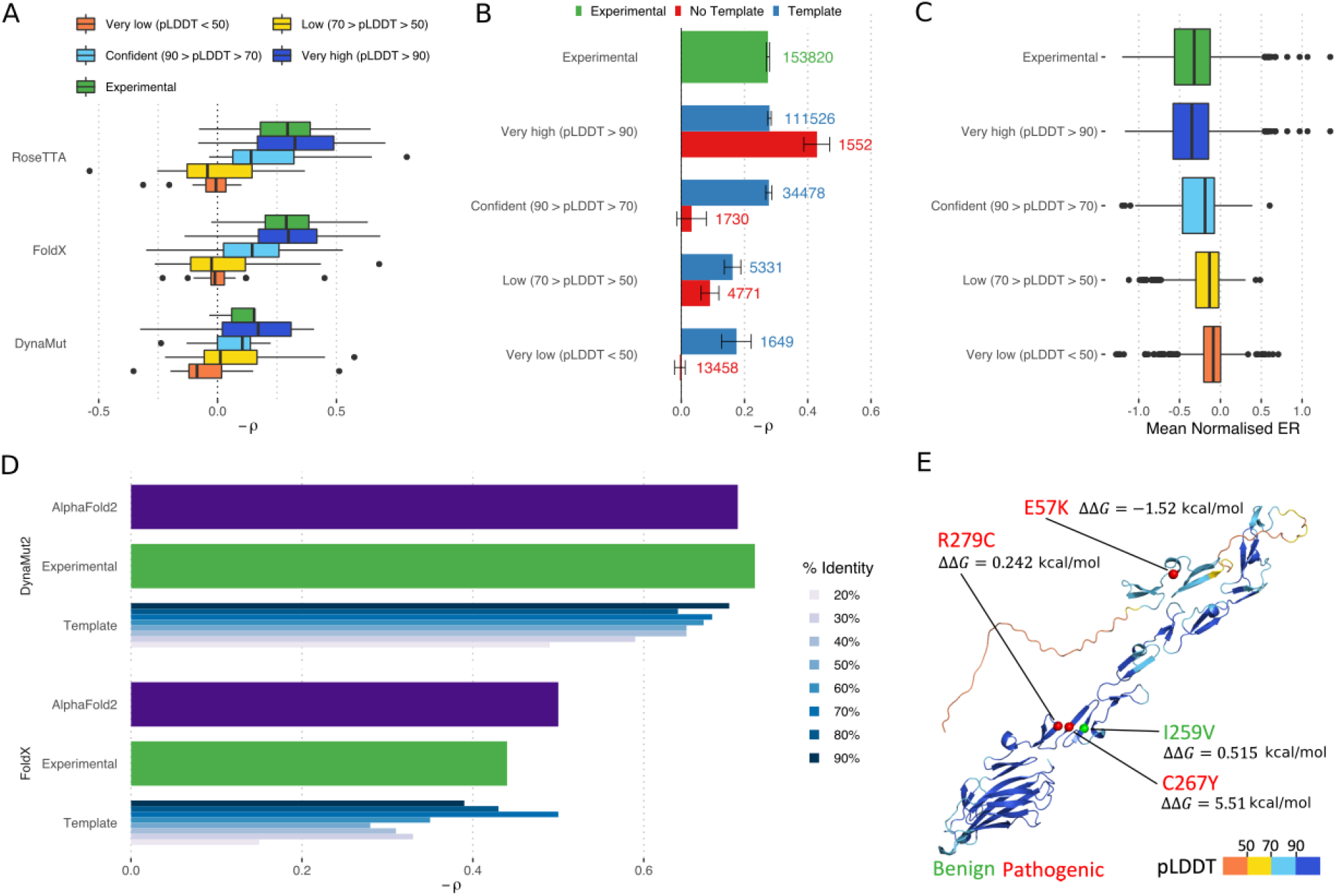
Comparing structure based prediction of impact of protein missense mutations using experimental or AF2 derived models. A) Relation between the predicted ΔΔG for mutations with measured experimental impact of the mutation from deep mutational scanning data (−1 * Pearson correlation). The predicted change in stability was done with one of 3 structure based methods using structures from AF2 or available experimental models. B) Correlations based on the FoldX predictions as in A) but subsetting the positions in AF2 models according to confidence and whether the position is present in an experimental structure. C) The mean impact of a mutation from DMS data for positions in AF2 models with different degrees of confidence. D) Comparative performance of methods predicting stability changes upon mutation using AF2, experimental and homology models based on temples of different identity cut-offs. Experimental measurements of stability are for 2,648 single-point missense mutations over 121 proteins. E) Example application for structure based prediction of stability impact of known disease mutations for a human protein with little structural coverage prior to AF2. ΔΔG stability changes were predicted using FoldX and a significant impact was considered for |ΔΔG|>1 kcal/mol.

The compilation of impact on protein stability contains information for 2,648 single-point missense mutations over 121 distinct proteins. We compared the accuracy of structure based prediction of stability changes using AF2 structures, experimental structures and homology model based predicted structures using different sequence identify cut-offs (Fig 3D and Fig S7, see Methods). Across 11 well-established structure based approaches to predict the impact of mutations (Fig 3D and Fig S7), the predictions based on AF2 models were comparable to experimental structures and homology model based predictions tended to show significant decreases in performance for templates below 40% sequence identity.

We investigated as an example the human EGF-containing fibulin-like extracellular matrix protein 2 (EFEMP2). EFEMP2 is a 443 residue long protein that previously had only a NMR model for 71 residues (2kl7). The AF2 prediction is largely of high confidence with the exception of the first 100 amino acids. As there are several disease-associated missense mutations annotated to this protein we predicted the impact of these on protein stability (Fig 3E) and obtained 3 out of 4 mutational impacts showing concordance to the mutation annotation as either pathogenic (predicted |ΔΔG|>1 Kcal/mol) or benign.

The systematic results and the specific example indicate that AF2 structures with confident predictions can be used to generate structural hypotheses about the potential impact of disease or trait-associated mutations.

### Functional characterization of AF2 models by pocket and structural motif prediction

High confidence proteome-wide structural predictions open the door for a large expansion of predicted protein pockets previously only possible for experimental structures or models based on close homologues (Bhagavat et al., 2018; Kana and Brylinski, 2019). However, the full protein models produced by AF2 have to be considered carefully given potential errors, such as the likely incorrect placement of protein segments of low confidence, or the low confidence in inter-domain orientations. To investigate if these issues may result in the formation of spurious pockets, we predicted pockets and bindings sites on a set of structures with known binding sites (UBS) defined using bound *(holo)* structures, and the corresponding unbound (*apo*) structures (Clark et al., 2020). We retained 230 of 304 protein families from this data set.

Pockets identified from structures have a wider size range than ground truth binding sites (Fig 4A). This is also true for pockets predicted from AF2 structures, which include a small number of particularly large pockets (Fig 4A). We averaged residue-level pLDDT values of the predicted pocket binding residues to define pocket-level estimates. These unusually large pockets show a lower mean pLDDT compared to other pockets (Fig 4B). We then asked whether mean pLDDT could be useful as a general metric of prediction confidence, regardless of the size of the pocket. We quantified the overlap between ground truth pockets (UBS) and pocket predictions, further splitting AF2 predicted pockets into high-quality (mean pLDDT >90) and low-quality (mean pLDDT ≤ 90) subsets (Fig 4C). Regardless of the overlap metric and pocket detection method used, we do not observe a difference between the performance of high-quality AF2 pockets, and pockets identified from experimental (apo or holo) structures. In contrast, low-confidence pockets do not generally overlap with ground truth binding sites. While there may be a bias in the fact that high confidence AF2 regions will more likely have relevant deposited templates, we suggest that the mean pLDDT of predicted pockets can be used as an additional criterion for pocket selection in AF2 structures.

**Figure 4.**
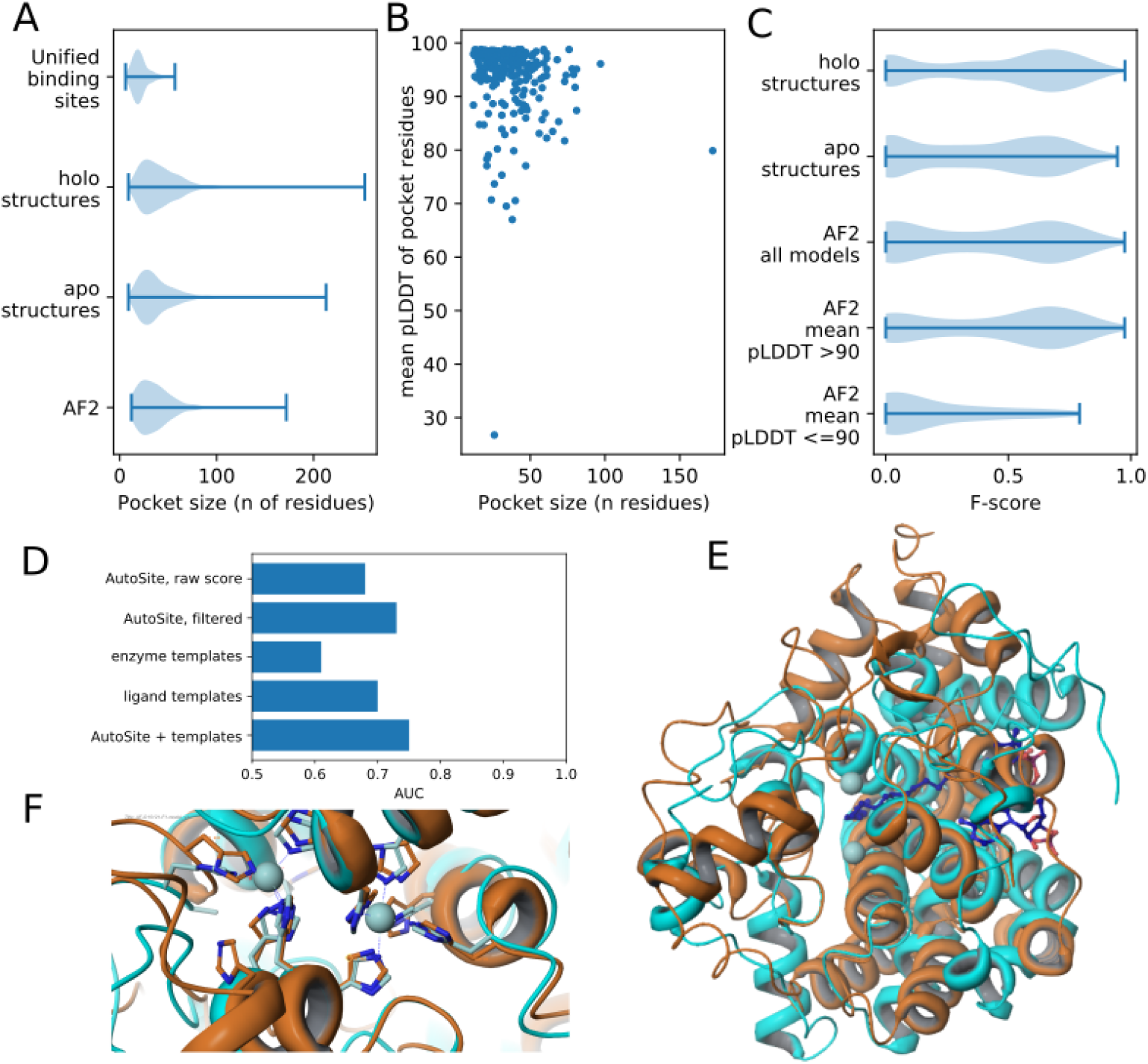
Pocket detection and function prediction. A) Size of known binding sites (UBS) compared to size of top AutoSite pockets in experimental holo (bound), experimental apo (unbound) and AF2 structures. B) Top pocket size versus mean pLDDT of pocket-associated residues. C) Distribution of overlaps between known binding sites and top predicted pockets for holo, apo and AF2 structures. AF2 structures are further subdivided by the mean pLDDT of pocket-associated residues.D) Enzymatic activity prediction using pocket-derived, template-derived, and combined metrics. E) Superposition of the AF2 model of DEGS1 (O15121) with 4zyo. Orange : ribbon representation of DEGS1 Cyan: ribbon representation of pdb structure 4zyo. Zinc atoms (light blue spheres) and bound substrate (dark blue ball and stick) as observed in the structure of 4zyo are also shown. F) Closeup around the metal binding centre of 4zyo. Ribbon representation of the protein and metal chelators for DEGS1 and 4zyo are shown in orange and cyan respectively. The zinc atoms observed in 4zyo are shown as light blue spheres. Metal chelating residues for DEGS1 are clearly identifiable.

Conserved local conformations of specific residues can identify important functions such as enzyme activity, ion or ligand binding beyond global sequence and fold similarities (Laskowski et al., 2005). To showcase the potential of such future application to AF2 models we focused on 100 human proteins with large contiguous regions of confident predictions with no models available in the Swiss Model repository. We combined residue template searches with pocket detection to predict which may have enzymatic activities. The combined pocket and template scores enriches the set for proteins with previous annotations for enzymatic activity (Fig 4D, Table S3). As a specific example we focused on the human Sphingolipid delta(4)-desaturase (EC 1.14.19.17, DEGS1, UniProt Accession O15121), which has a high confidence level (average pLDDT=96.31) with no previous structural data. A sequence search of the 323 residue protein against all existing entries in the PDB database shows that the best sequence match is 23.5%, with PDB entry 1vhb (Bacterial dimeric hemoglobin, 9115439), indicating the lack of any structural models from homology. Scanning 400 auto-generated 3-residue templates from the AlphaFold predicted structure against representative structures in the PDB (reverse template comparison, (Laskowski et al., 2005)) reported a possible three-residue template match with PDB entry 4zyo (EC 1.14.19.1, human stearoyl-CoA desaturase (Wang et al., 2015), Fig 4E). A closeup of the metal binding centre (Fig 4F) of DEGS1 and 4yzo (overall sequence homology 12.1%) superimposed via the three residue templates (Fig 4E) clearly indicates the potential dimetal catalytic center for DEGS1. The histidine-coordinating metal centre of DEGS1 thus identified, together with data on the bound substrate of 4zyo, provides a foundation for modelling studies to impact on the pharmacology of DEGS1 by exploring the details of its catalytic mechanism.

These results illustrate the large potential for AF2 models to be applied on a large proteome-wide scale for the discovery of potential protein function, characterization of under-explored drug binding sites and characterization of enzymatic potential across species.

### AF2 based prediction of protein complex structures

Since the first development of direct coupling analysis algorithms, co-evolutionary information based methods have been used to predict protein-protein interactions (Weigt et al., 2009). Lately, several deep learning-based methods such as trRosetta (Yang et al., 2020) and Raptor-X (Jing et al., 2020) have been reported to be able to predict the structure of some protein complexes. While AF2 was trained to predict the structure of individual chains, it is possible that it can also be used to predict the structure of protein complexes, just as trRosetta, Raptor-X and RoseTTAFold. To examine this, we tested the ability of AF2 to fold and “dock” two benchmark sets — a set of proteins known to form oligomers (Ponstingl et al., 2000) and the Dockground 4.3 heterodimeric benchmark (Kundrotas et al., 2020).

For oligomerization we obtained sets of proteins known either not to oligomerise or to form oligomers including dimers, trimers or tetramers. We then made AF2 predictions for each protein attempting to predict either a monomer or an oligomeric form (see Methods). Across the set of predictions higher scores are given to models corresponding to the correct oligomerization state with 71 out of 87 (82%) predicted top scoring models corresponding to the correct state (Fig 5A, Table S4) and generally the multimeric state scores are well separated from the monomeric state scores (Fig 5B). In 28/30 examples, AF2 was able to correctly predict monomeric proteins as monomers, 29/35 dimers as dimers, 7/9 trimers as trimers, and 7/13 tetramers as tetramers. Interestingly, even though the failure rate is high for tetramer state predictions, the predicted structure for the corresponding state was actually correct for 5/6 failures. Examples of failure modes for dimers and a tetramer are shown in figures Fig 5C and Fig 5D. We noted that for some cases of failed tetramer predictions we could obtain higher confidence of the tetramer predictions by increasing the number of recycles.

**Figure 5.**
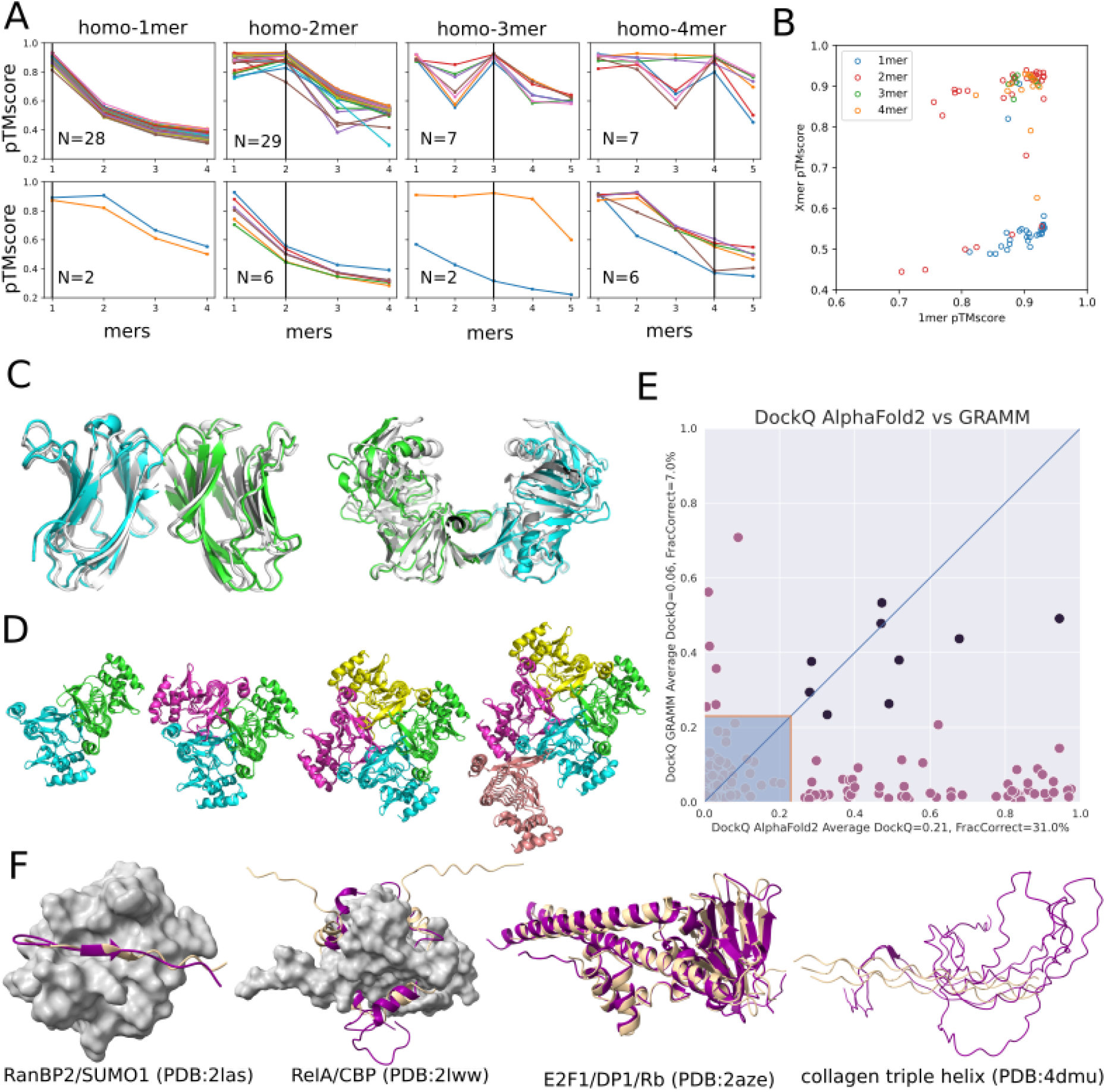
Using AF2 to predict homo-oligomeric assemblies and their oligomeric state. A) AF2 prediction for each oligomeric state (1–4 for monomers and dimers, and 1–5 for trimers and tetramers). Only proteins where the monomer had predicted LDDT over 90 are shown. For visualization, the predicted successes (top) and failures (bottom) were separated into two different plots. Success is defined when the peak of the homo-oligomeric state scan matches the annotation or the pTMscore of the next oligomer state is significantly lower (−0.1). B) For each of the annotated assemblies the pTMscore of monomeric prediction is compared to the max pTMscore of non-monomeric prediction. C) Monomer prediction failure. Two monomers were predicted as homo-dimers. For the first case (pdb:1bkz), the prediction matched the asymmetric unit (shown as blue/green and prediction in white). For the second case (pdb:1bwz), the prediction matched one of the crystallographic interfaces. D) 3TDT trimer predicted as tetramer. Though the interface is technically correct, for this c-symmetric protein, pTMscore is not able to discriminate between 3 and 4 copies. E) Comparison of docking quality between AF2 (X-axis) and a standard docking tool GRAMM (Y-axis) (Tovchigrechko et al., 2002). Comparisons are made using the DockQ score (Basu and Wallner, 2016; Tovchigrechko et al., 2002). Models with a DockQ score higher than 0.23 are assumed to be acceptable according to the CAPRI criteria (marked outside the shaded area). Black circles indicate the complex was well modelled with both methods. The average DockQ score and the number of acceptable or better models are shown in the axis labels. It should be noted here that AF2AlphaFold2 both folds and docks the proteins, while GRAMM only docks them. F) Examples of AF2 predicted interactions mediated by regions of intrinsic disorder.

We next examined the Dockground 4.3 heterodimeric benchmark set (Kundrotas et al., 2020). We predicted complex structures using the DeepMind default dataset and the small BFD database. It can be noted here that this method does not include any “pairing”of interacting chains as used in earlier Fold and Dock approaches. The docking quality was evaluated using DockQ (Basu and Wallner, 2016; Tovchigrechko et al., 2002). Only one model for each target was made and a maximum of three recycles were allowed. In Fig 5E it can be seen that the performance is far superior to traditional docking methods, with 31% of well corrected predicted protein complex models compared with 7% using GRAMM, a standard shape complementarity docking method (Tovchigrechko et al., 2002).

Finally, we studied examples of complexes containing intrinsically disordered proteins/regions (IDPs/IDRs) that adopt a stable structure upon binding. IDRs often bind via short linear motifs (SLiMs) recognizing folded domains driven by a few residues. The longer IDRs can contain arrays of SLiMs or can also form stable structures upon binding to other IDRs without a structured template. We selected 14 cases of complexes involving IDRs with known structures and analysed in detail the distinguishing features compared to the experimental complex (Fig 5F for selected examples and SFig 8 and SFig 9 for all examples). In general, AF2 performs remarkably well in predicting SLiMs fitting into a well-defined binding pocket driven by hydrophobic interactions, such as the SUMO interacting motif of RanBP2. Longer IDRs, often containing tandem motifs, are often challenging, especially if they have a symmetric structure. For the RelA:CBP interaction, AF2 correctly finds the binding groove but fits the IDR in a reverse orientation. AF2 also performs well on complexes where IDRs are part of a multi-IDR single folding unit, such as the E2F1-DP1-Rb trimer, however, building complexes for proteins with highly unusual residue compositions, such as collagen triple helices, often fail. We provide a detailed description of the 14 examples in SFig 8 and SFig 9 and Table S5 detailing the factors that enable or hinder successful predictions.

For complexes, AF2 demonstrates an accuracy in predicting the structure of homo and hetero oligomers for both folded and disordered binding partners beyond other current state-of-the art methodologies. This is an unintended application of AF2 that may be further improved when protein interaction specific training is implemented.

### Evaluation of AF2 models for use in experimental model building

The remarkable accuracy of AF2 predictions even in the face of little to no sequence similarityhomology to existing structures provides clear opportunities for future use in experimental model building: (a) the use of AF2 models for molecular replacement or docking into cryo-EM density in place of homologous structures, experimental phasing and/or *ab initio* model building; and (b) use of AF2 models as reference points to improve existing low-resolution structures. These use cases will typically involve the use of conformational restraints, e.g. to maintain the local geometry of domains while flexibly fitting a large multi-domain model, or to restrain the local geometry of an existing model of an AF2-derived reference to highlight and correct likely sites of error. It is critical to use restraint schemes designed to avoid forcing the model into conformations that clearly disagree with the data. Typically, this is achieved via some form of top-out restraint, for which the applied bias drops off at large deviations from the target. Here, we take advantage of the fact that AF2 models include typically very strong predictions of their own local uncertainty to adjust per-restraint weighting of the adaptive restraints recently implemented in ISOLDE (Croll, 2018) (see Methods). For the two case studies discussed below, a comparison of validation statistics for the original and revised models may be found in Table S6.

As an example of the improvement of existing structures we used the eukaryotic translation initiation factor (eIF) 2B bound to eIF2 (6o85), a 0.4 MDa 8-chain complex captured by cryo-EM at an overall resolution of 3 Å (Kenner et al., 2019). Rigid-body alignment of AF2 models to their corresponding experimental chains (Fig 6A) showed overall excellent agreement, with the largest deviations corresponding to correctly-folded domains with flexible connections to their neighbours. Other mismatched smaller regions correspond to either register errors in the original model or flexible loops and tails. Each chain was restrained to its corresponding AF2 model using ISOLDE’s reference-model distance and torsion restraints, with each distance restraint adjusted according to pLDDT. Future work will explore the use of the predicted aligned error (PAE) matrix for this purpose, and weighting of torsion restraints according to pLDDT.

**Figure 6:**
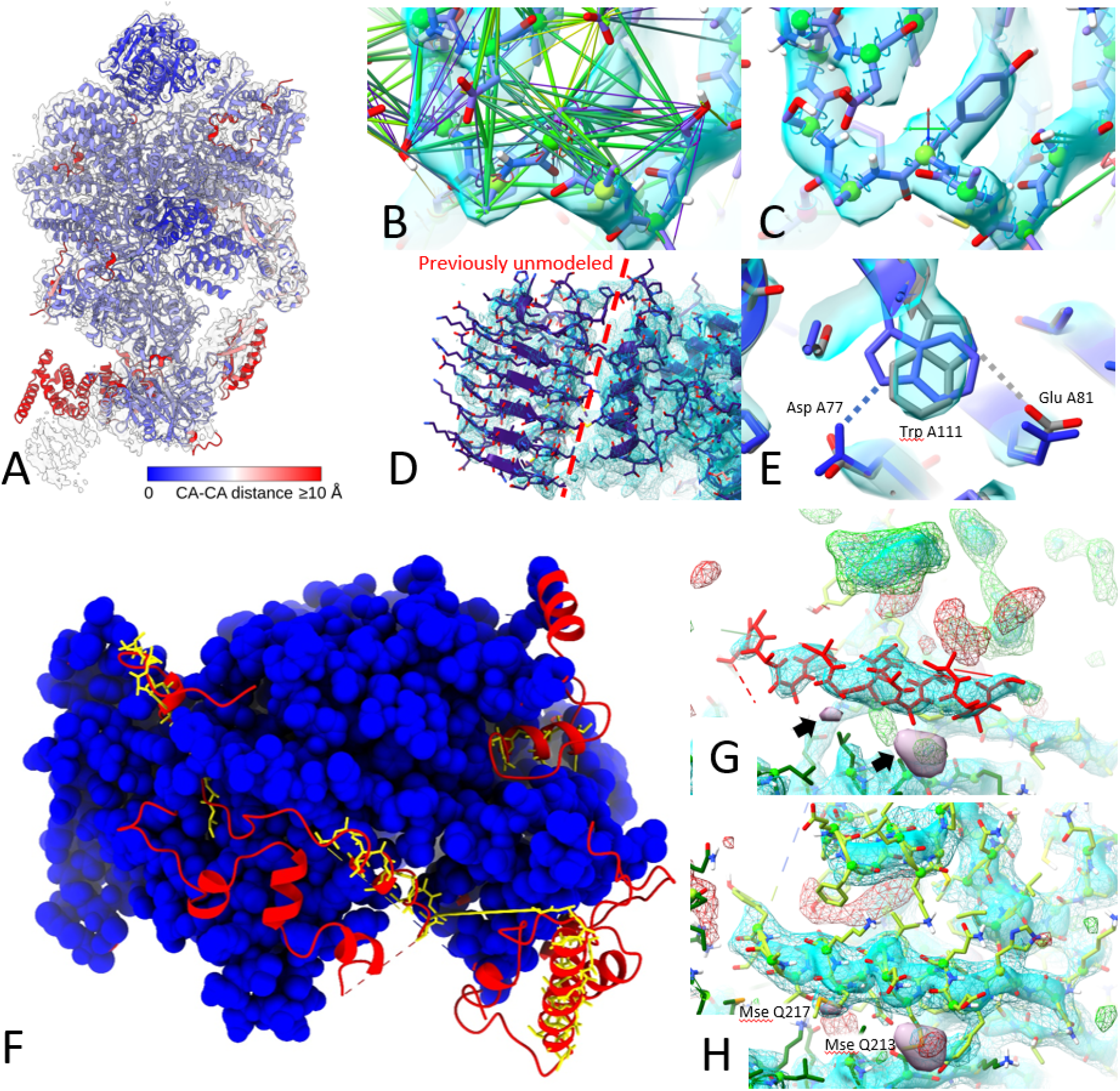
Application of AF2 predictions to modelling into cryo-EM (A-E) or crystallographic (F-H) data. A) AlphaFold2 predictions for individual chains in 6o85, aligned to the original model and colored by Cα-Cα distance, with the map (EMD-0651) contoured at 6.5 σ. Red domains at bottom were correctly folded but misplaced due to flexibility; smaller regions of red correspond either to flexible tails or register errors in the original model. (B-C) use of adaptive distance and torsion restraints to correct problematic geometry in the original model. B) before; C) after refitting; satisfied distance restraints are hidden for clarity. D) Due to very poor local resolution and lack of homologues, the C-terminal domain in chain J (left of the dashed line) was previously left unmodelled. This domain was predicted with high confidence by AlphaFold2 (mean pLDDT=83.0), and fit readily into the available density. E) High-confidence regions may still contain subtle errors that are difficult or impossible to detect in the absence of experimental data. The sidechain of Trp A111 (pLDDT=86.1) was modelled backwards (blue), forming a H-bond to Asp A77; the final model fitted to the map (grey) instead H-bonds to Glu A81. F) Rebuilding the recent 3.3 Å crystal structure 7ogg starting from molecular replacement with AlphaFold2 models dramatically improved model completeness. Blue spacefill = residues identified in original model; yellow sticks = residues modelled as unknown (UNK) in original model; red cartoon: residues identified in rebuilt model. G) Helix modelled as UNK (residues 558-573 of chain R, red), surrounded by unmodelled density (3 σ mFo-DFc, green(+), red(-); +2 σ sharpened 2mFo-DFc, cyan surface; +1.5 σ unsharpened 2mFo-DFc, cyan wireframe; +5 σ anomalous difference map, purple surface and arrows). H) Final model, with anomalous difference blobs corresponding to selenomethionine residues 213 and 217 of chain Q and the previously unmodelled density filled; this region was predicted with an average pLDDT of 88, and required only minor sidechain corrections to fit the density.

Simple energy minimisation and equilibration of the restrained model at 20 K corrected the majority of local geometry issues (e.g. Fig 6B and 6C); a high-confidence prediction for the C-terminal domain of chains I and J allowed us to add this into previously-untraceable low resolution density (Fig 6D, left of the dashed line). We emphasize that detailed manual inspection remains necessary to find and correct larger errors in the experimental model, sites of disagreement arising from conformational variability, and sites where high-confidence predictions are in fact incorrect. An example of the latter is the sidechain of Trp A111, which despite its high confidence (pLDDT=86.1) was modelled incorrectly by AF2 (Fig 6F).

To explore the use of AF2 structures for solving and refining new structures, and to map out suitable workflows, we attempted to recapitulate the recent 3.3 Å crystal structure of the *S. cerevisiae* Nse5/6 complex (7ogg) (Taschner et al., 2021). This was not included in the AF2 training set and no existing structures have ≥30% identity to either chain. Originally solved via selenomethionine experimental phasing, the combination of low resolution and anisotropy (ΔB = 80 Å^2^) meant that only 600 out of 850 residues were definitively modelled by the authors, with a further 65 residues traced as unknown sequence. For testing purposes we discarded this model and used the AF2 predictions for molecular replacement (MR). MR requires very close correspondence between atom positions in the search model and in the crystal; separation into individual rigid domains and trimming of flexible loops is a necessity. Here we used the PAE matrix to extract a single rigid core from each chain (see Methods) and performed MR in Phaser (McCoy et al., 2007) leading to a clear solution with TFZ=28.2, LLG=884 (see Methods).

Currently, a refined MR solution is typically used as the starting point for some combination of automatic and manual building of missing portions into the density. In many cases, however, it appears that AF2 predictions will support a more “top-down” approach, in which all residues predicted with at least moderate confidence are present in the initial model. To explore this, we trimmed the predicted chains to exclude residues with pLDDT ≤ 50 and aligned the result to the MR solution, setting the occupancies of all atoms not used for MR to zero. This was used as the starting point for rebuilding in ISOLDE; here zero-occupancy atoms do not contribute to structure factor calculations or bulk solvent masking, but still take part in molecular interactions and are attracted into the map. The model was subjected to three rounds of end-to-end inspection and rebuilding interspersed with refinement with phenix.refine (Liebschner et al.,2019). In the initial round, zero-occupancy residues fitting the map were reinstated to full occupancy, while residues which appeared truly unresolved were deleted; a small number of these were re-introduced in subsequent rounds. The total time spent was approximately one working day; the final model (Fig 6 F-H) increased the number of modelled, identified residues from 600 to 818, while slightly improving overall geometry and reducing the R_free_ from 0.317 to 0.295. With few exceptions (primarily at heterodimer and symmetry interfaces), rebuilding was limited to minor sidechain adjustments. Thus, in this case the availability of high-quality structure predictions has reduced an extremely challenging, near-intractable structure solution into a straightforward task.

## Discussion

We have estimated that AlphaFold 2 (AF2) may add, on average, around 25% of confidently predicted residues to a given proteome, although this will vary depending on how much experimental and previous approaches could already cover. However, even for residues that can be modelled by distant homology it is possible that the AF2 models are more accurate, increasing the usefulness of these models. Here the very precise estimates of accuracy at the residues level are extremely useful for a user. Given that AF2 depends on multiple sequence alignments that are often easier to obtain for bacterial proteins, these species will also tend to have a higher proportion of the proteome possibly covered by high confidence predictions. Across the 11 species we could compare with the SwissModel repository, *P. falciparum* remains perhaps the most challenging to model, likely due to sparsity of homologous sequences/structures. Sequencing efforts in metagenomics or more broadly aimed at covering the tree of life will improve the capacities of AF2 and related approaches. In addition, low AF2 scores are highly enriched for protein disorder and metrics derived from AF2 predictions outcompete state-of-the-art disorder predictions. This suggests that regions of low confidence predictions can be hypothesized as disordered segments.

The AF2 database was initially released with >300k proteins modelled and is planned to scale to over 100 million, sampling the universe of protein sequences and structures. As we show here, even on a relatively modest set, we can identify what are likely to correspond to rarely explored structural elements. Among other areas, the expansion of high confidence predictions will allow for the prioritization of experimental structure determination of novel folds; the large scale prediction of protein function from structure; the identification of novel enzymes; and the study of evolution of protein folds.

We have assessed the application of AF2 predictions in diverse structural biology challenges including variant effect prediction, pocket detection and model building into experimental data. In line with the reported high accuracy of the models, we find that AF2 predicted structures on average tend to give results that are as good as those derived from experimental structures. However, while AF2 returns full protein predictions, these can often contain protein segments that are placed with uncertainty. For example, AF2 is unaware of where transmembrane regions are and can be uncertain about the relative orientation of different domains within a protein. This uncertainty can lead to incorrect estimations or identification of folds, pockets, variant effects or poor model building. Importantly, in all cases we find that it is critical to take into account the confidence metrics provided and that these should be incorporated into the corresponding workflows. Taking these into account leads to improvements in all of the tasks that we have assessed here.

Finally, we explored the application of AF2 for prediction of complex structures finding that it outcompetes standard docking approaches while at the same time not requiring even starting protein structures. We have expanded on this analysis in a companion manuscript (Bryant et al.,2021). It has already been shown that other distance/contact-prediction methods trained to predict intra-chain contacts can be used unmodified to predict inter-chain contact predictions, both for homo- and hetero-meric complexes. Therefore, we were not surprised to see that it is possible to use AlphaFold 2 to Fold and Dock heterodimeric complexes. However, it was unexpected that it was possible to use non-matched pairs of alignments for different proteins to predict complex structures using AF2 indicating that AF2 goes beyond using co-evolution to predict these structures. It is possible that approaches specifically trained or fine-tuned to predict complexes may outperform AF2 even if training such methods may be challenging given the smaller datasets of protein complexes compared to individual proteins. In any case, the promising results of AF2 shows that accurate predictions of protein-protein interactions may likely be achieved for most pairs.

In summary, we find that AF2 models, when considering their uncertainty, can be applied to existing structural biology challenges with near experimental quality. The application of AF2 to a large representation of the protein universe and expansion to the prediction of protein complexes will have a transformative impact in life sciences.

## Methods

### Coverage comparison between SwissModel repository and AlphaFold 2 databases

The SwissModel repository (SMR) and AlphaFold 2 (AF2) databases were accessed on 24.07.2021. Reference proteomes for 11 species common to AF2 and SMR were downloaded from the Uniprot release 2021_03. Only structures corresponding to entries from the reference proteomes were used for the analysis. Numpy (Harris et al., 2020), Pandas (Reback et al.,2021), Prody (Bakan et al., 2011), and Matplotlib (Caswell et al., 2021) Python libraries were used for the analysis and the visualization. Structure counters for protein domains were extracted from the corresponding InterPro entries (Blum et al., 2021). Code and data are available online (https://github.com/aozalevsky/alphafold2_vs_swissmodel/).

### Comparison between human RoseTTAFold Pfam domain and AF2 structures

We used the 17,006 human proteins that were described as the principal isoform for their corresponding gene according to APPRIS and whose sequences were the same in ENSEMBL and Uniprot. We used Pfamscan to identify PFAM domains in the 17,006 protein sequences. The database of PFAM-A models was downloaded on June 29th 2021 and created on March 19th 2021. We only kept those PFAM domains identified with an e-value below 1e-8. AF2 models for human proteins were downloaded on July 23rd 2021 from https://alphafold.ebi.ac.uk. We extracted the sequences and compared them to the ENSEMBL protein sequences used for the structural analysis. For comparison purposes all the analyses and results presented here are based on the subset of 17.006 protein sequences for which the ENSEMBL and AlphaFold protein sequences were identical. We also extracted pLDDT values for each residue from the AlphaFold models, as these are stored as if they were the B-factor of the protein coordinates file. The RoseTTAFold models were downloaded from the EBI website on July 27th 2021 and the RMSD between models from both methods was calculated using the function *struct.aln* from the R package “bio3d”. All statistical analyses were done using R 4.0.2. Graphical plots were created with the packages “ggplot2” (Wilkinson, 2011), “patchwork” and “reshape2”. Molecular graphics and analyses performed with UCSF Chimera (Pettersen et al., 2004), developed by the Resource for Biocomputing, Visualization, and Informatics at the University of California, San Francisco, with support from NIH P41-GM10331.

### Disorder prediction

Benchmarking data for ordered and disordered protein regions were taken from the benchmark set of IUPred2 (Mészáros et al., 2018) filtering for proteins for which AF2 predicted structures are available in the AlphaFold database. Relative solvent accessible surface area (relative SASA) was calculated by determining the absolute SASA using DSSP and then comparing it to the SASA calculated in a Gly-Gly-X-Gly-Gly conformation. ROC curves plotting the true positive rate as the function of the false positive rate were calculated on a per-residue level. Area under the ROC curves (AUC) are single number measures of the overall predictive performance in the range of 0.5 (for random predictions) to 1.0 (for perfect predictions).

### Exploration of structural space covered by AlphaFold DB compared to the PDB

We use the 365,198 proteins from the current AlphaFold database (AF) and 104,323 proteins from PDB in 2016 (until CASP12) with a 100% sequence identity threshold, removing duplicates. Due to the presence of low confidence regions in the AF proteins we first perform trimming to split each AF protein into smaller high confidence units as follows: A 1D Gaussian filter with a σ of 5 is applied to the sequence of pLDDT scores extracted from the Cα atoms. The resulting scores are used to split the protein into continuous segments of residues with smoothed pLDDT scores > 70. Segments with less than 50 residues are discarded. This removed 68,890 AF proteins with too few high confidence residues for accurate structural comparison.

For each AF protein segment, and for each PDB protein, rotation invariant moments O3, O4, O5, and F (as described in Geometricus (Durairaj et al., 2020)) are calculated for the Cα atoms using a k-mer based approach (with k=16) and radius-based approach (r=10 Å). These are then converted into shape-mers using a resolution of 4 for the k-mer based approach and 6 for the radius-based approach. Shape-mers are counted across the whole protein for a PDB protein and across all splits for an AF protein to give the shape-mer count vectors. We then created a term frequency inverse document frequency (TFIDF) matrix for all PDB and AF proteins, where the terms are shape-mers and each protein is equivalent to a document. We performed topic modelling using Non-negative Matrix Factorization (NMF), which attempts to factorize a matrix of size n×m into matrices W of size n×p, and H of size p×m. We interpret this as finding p topics (here set to 250), each of which consists of a weighted combination of the m shape-mers (defined by H). Each of the n proteins can then be seen as a weighted combination of these p topics (defined by W).

For topic analysis, we assigned proteins to each topic using knee detection with a weight cut-off (Satopaa, Ville, et al. 2011). For visualization, we performed t-SNE dimensionality reduction on the W matrix returned by NMF. Topic-specific scores were obtained for each residue within a shape-mer by multiplying the corresponding topic weight for the shape-mer (from H) with an RBF kernel score of the Euclidean distance between the residue and the central residue of the shape-mer. These were aggregated across all shape-mers within a protein to obtain the topic-specific residue scores for the protein.

Code and scripts can be found at https://github.com/TurtleTools/alphafold-structural-space

### Structure based variant effect predictions using experimental and predicted structures

A subset of experimentally characterised mutations was curated from ThermoMutDB (Xavier et al., 2021), comprising 2,648 single point missense mutations across 132 unique globular proteins. The experimentally measured effect of the mutations on protein stability was represented as the difference in Gibbs Free Energy (**ΔΔ**G in kcal/mol) between wild-type (**Δ**GWT) and mutant (**Δ**GMT). Experimental structures were obtained from the PDB (Burley et al., 2021). Homology models were generated using Modeller (Sali and Blundell, 1993) using the most complete available template within each identity threshold range (20% ± 5%, 30%± 5%, until 90% ± 5%). AF2 models were generated locally. These mutations were analysed by computational predictive tools including the sequence based predictors I-Mutant (Capriotti et al.,2005), SAAFEC-SEQ (Li et al., 2021), MUpro (Cheng et al., 2006), and the structure based predictors mCSM-stability (Pires et al., 2014a), DUET (Pires et al., 2014b), SDM (Worth et al.,2011), DynaMut (Rodrigues et al., 2018), MAESTRO (Laimer et al., 2015), ENCoM (Frappier et al., 2015), DynaMut2 (Rodrigues et al., 2021), FoldX (Delgado et al., 2019). Methods’ performance and concordance between the experimental and predicted **ΔΔ**G are presented as Pearson’s correlations. A larger set of experimentally determined impact of missense mutations was derived from a compilation of Deep Mutational Scanning (DMS) experiments (Dunham and Beltrao, 2021) comprising 117,135 total mutations in 33 proteins. These were compared against predictions made with DynaMut2, FoldX and Rosetta, calculated as previously described (Høie et al., 2021; Park et al., 2016) (see also suppl. Methods).

### Pocket and structural motif prediction

We downloaded structures from the AF2 public database (https://alphafold.ebi.ac.uk) except for analyses of the LBSp dataset (Clark et al., 2020). In the latter case, we used locally modelled structures, as many LBSp structures are from species not included in the public database. We detected pockets and calculated overlap metrics (F-score, MCC) as described in (Clark et al.,2020). As an exception, we used OpenBabel (O’Boyle et al., 2011) to prepare PDBQT input for AutoSite (obabel -h -xr --partialcharge gasteiger). We used AutoSite (Ravindranath and Sanner,2016) from ADFRsuite version 1.0 and ghecom (Kawabata, 2010) release 2018/06/15. We considered a protein as having known enzymatic activity if there was an EC number and/or a catalytic activity annotation in its UniProt record.

### Oligomerization state prediction

To test the ability of AlphaFold to predict the oligomeric state of homo-oligomeric assemblies, we downloaded the dataset described in Ponstingl et al (Ponstingl et al., 2000). Since the PDB files were not provided, the dataset was filtered to entries where the oligomeric state was in agreement with PISA annotation. Since AlphaFold’s training was only done on single chains, we reasoned that examples, even if they overlap with the training set, could be used to evaluate AlphaFold’s oligomeric state prediction capabilities. For each PDB, the sequence of chain A was extracted and a multiple sequence alignment was generated using the automated MMseqs2 webserver through ColabFold. For homo-oligomeric prediction, each MSA was copied, padded with gaps to the total length reflecting the number of copies in the assembly, and concatenated. These concatenated alignments were fed into AlphaFold. No templates were used. All five ptm-fine tuned model parameters were used. To test the robustness of AF2’s five model parameters to predict homo-oligomeric structures, we use the worst of the predicted TMscores for each state.

### *Fold and Dock prediction of* heterodimeric *protein complexes*

We used 219 heterodimeric complexes from Dockground benchmark 4 (Kundrotas et al., 2018). This set contains unbound forms of heterodimeric protein chains, which share at least 97% sequence identity with the bound forms. The dataset consists of 54% eukaryotic proteins, 38% bacterial and 8% from mixed kingdoms, e.g. one bacterial protein interacting with one eukaryotic. To evaluate performance, one model for each pair was generated with AF2 (using default parameters, except that model_2 was used, providing a complementary set of results to those derived in (Bryant et al., 2021)). To enable docking we only changed the residue number so that both chains are treated as a long chain with a 200 residue gap, as described elsewhere (Mirdita et al.). For comparison models docked with GRAMM were used. The GRAMM models were ranked using the AACE18 scoring function (Anishchenko et al., 2018). Docking quality was estimated with DockQ (Basu and Wallner, 2016).

### Complex structure predictions for disordered proteins

Predictions were run using the sequences defined in the PDB files (not including modified residues and other molecules). Predictions were done using the Google Colab notebooks by Sergey Ovchinnikov; homooligomers were predicted using the notebook accessible at https://colab.research.google.com/github/sokrypton/ColabFold/blob/main/AlphaFold2.ipynb and heterooligomers were predicted using the dedicated notebook, accessible at https://colab.research.google.com/github/sokrypton/ColabFold/blob/main/AlphaFold2_complexes.ipynb. In the case of dimers, the default settings were used. In case of higher order oligomers, one chain was used on its own (usually the IDR if there is only one), and the rest of the chains were concatenated using a long linker (either several ‘U’s or several repeats of ‘SG’s).

### Evaluation of AF2 models for use in experimental model building

AF2 models were used as an aid to rebuilding the existing 6o85 in ISOLDE, with a preliminary implementation pLDDT-based weighting of its existing adaptive distance restraints (Croll, 2018). Initial fetching and alignment of the relevant AF2 models for each chain used a tool available in pre-release versions of ChimeraX 1.3, allowing command-based fetching of predictions from the AlphaFold EBI server by UniProt ID. For existing models fetched from the wwPDB, the UniProt ID for each chain is automatically parsed from the mmCIF metadata, and each fetched prediction is aligned and renamed to match the target chain. ISOLDE’s reference-model distance restraint scheme has four adjustable parameters controlling the restraint potential: kappa (the overall strength); wellHalfWidth (the range over which the restraining force is ~linearly related to distance); tolerance (the width of a flat-bottom - i.e. zero-force - region close to the target); and fallOff (the rate at which the potential tapers at large distances). With the exception of kappa each of these terms is expressed as a function of the reference interatomic distance: for a given restraint, the final harmonic well width, tolerance and fall-off all increase with increasing reference distance. For the purpose of this study, we added terms to further adjust kappa, tolerance and fallOff according to the pLDDT of the lowest-confidence atom in each restrained pair; all restraints where at least one reference atom had a pLDDT < 50 were disabled. For each chain in the complex, the working model was restrained against the AlphaFold reference using the “isolde restrain distances” command with the above modifications enabled but otherwise standard settings. Backbone and sidechain torsions were also restrained against the reference model using the “isolde restrain torsions” command with default arguments. After energy minimization and equilibration, the model was inspected and, where necessary, interactively remodelled; where reference model restraints clearly disagreed with the model, they were selectively released. Where the AF2 models included previously-unmodelled residues supported by the density, they were merged into the working model. The final model was refined with phenix.real_space_refine (Liebschner et al., 2019) using settings defined by the “isolde write phenixRsr” command.

For the recapitulation of 7ogg, the AF2 predictions for its two chains were fetched in ChimeraX as above. The rigid core of each chain was extracted using a community clustering approach based on the PAE matrix; source code for this is available at https://github.com/tristanic/pae_to_domains. After setting B-factors to a constant value of 50, these were used to generate a fresh molecular replacement (MR) solution using PHASER (McCoy et al., 2007). The original, complete AF2 models were aligned to the MR result in ChimeraX, and occupancies for atoms not part of the MR models were set to zero. The result was used as the starting model for rebuilding in ISOLDE. After initially settling into the map with distance and torsion restraints applied, the model was inspected and rebuilt end-to-end. During this initial rebuilding, zero-occupancy atoms with clear correspondence to density were reinstated to full occupancy, while residues with no associated density were deleted. Where there was clear disagreement with the map (primarily at the heterodimer interface), the initial distance and torsion restraints were selectively released in favour of interactive remodelling. The resulting model was refined with phenix.refine [ref] using settings defined by the “isolde write phenixRefine” command. In a second and third round of interactive rebuilding in ISOLDE (during which the distance and torsion restraints were fully released) interspersed with phenix.refine, a small number of residues deleted in the first step were re-introduced.

In both the above cases, the final coordinates have been shared with the original authors.

## Supporting information

SDataset 1

SDataset 2

## Supplementary Figures

**Fig S1.**
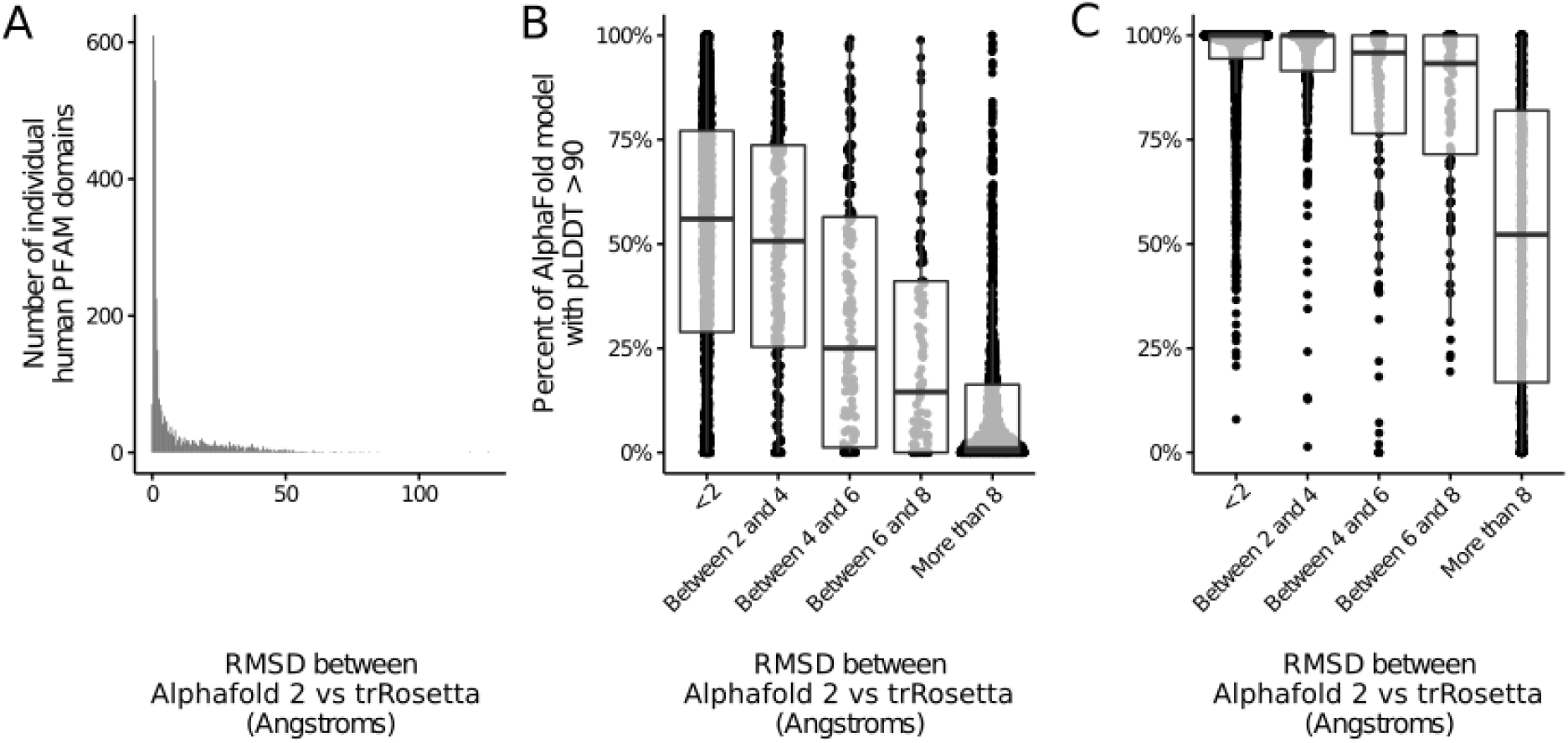
Comparison between AF2 and trRosetta. A) Histogram showing how many AlphaFold models for human PFAM domains (y-axis) depending on their RMSD to the generic RoseTTAFold model for the same PFAM domain (x-axis). B) Boxplots showing, for each of the 3035 AlphaFold models of human PFAM domains (each dot) the percent of residues with a pLDDT above 90 (y-axis). AlphaFold models are grouped by their RMSD to the trRosetta generic PFAM model (x-axis). C) Same as B but for residues with pLDDT higher than 50

**Fig S2.**
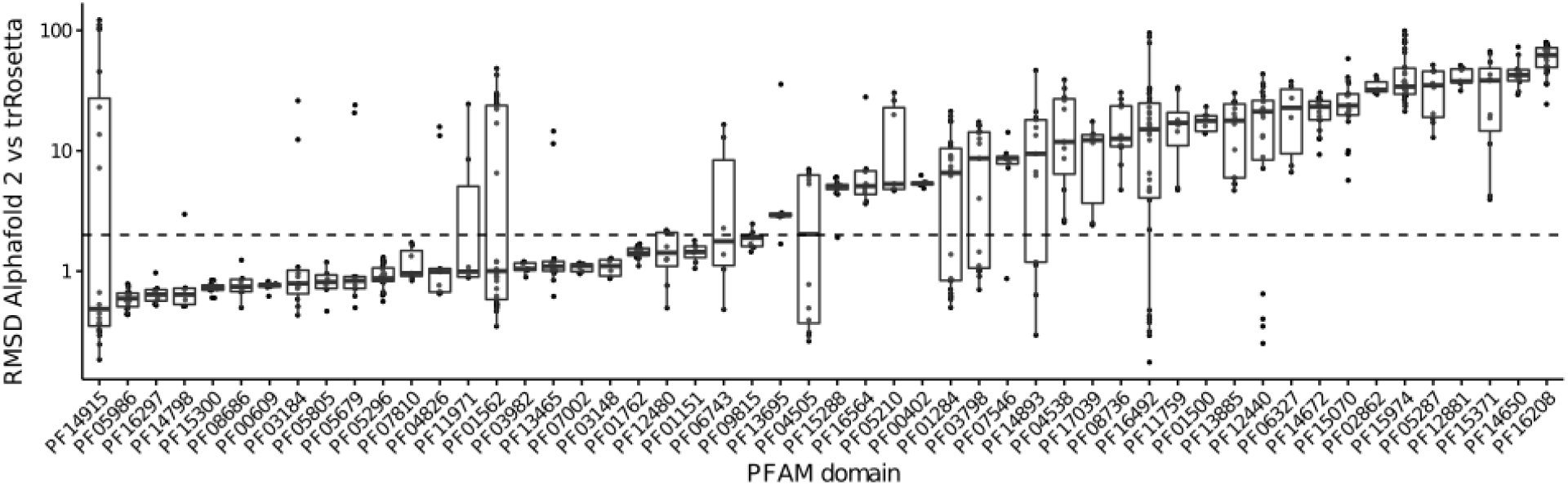
Intradomain variability between the predicted models of each tool. Boxplots showing the RMSD between AlphaFold and trRosetta (y-axis, logarithmic) for different PFAM domain instances in the human proteome (dots). Instances are grouped by PFAM families (x-axis). The black dashed line indicates an RMSD of 2Å.

**Fig S3.**
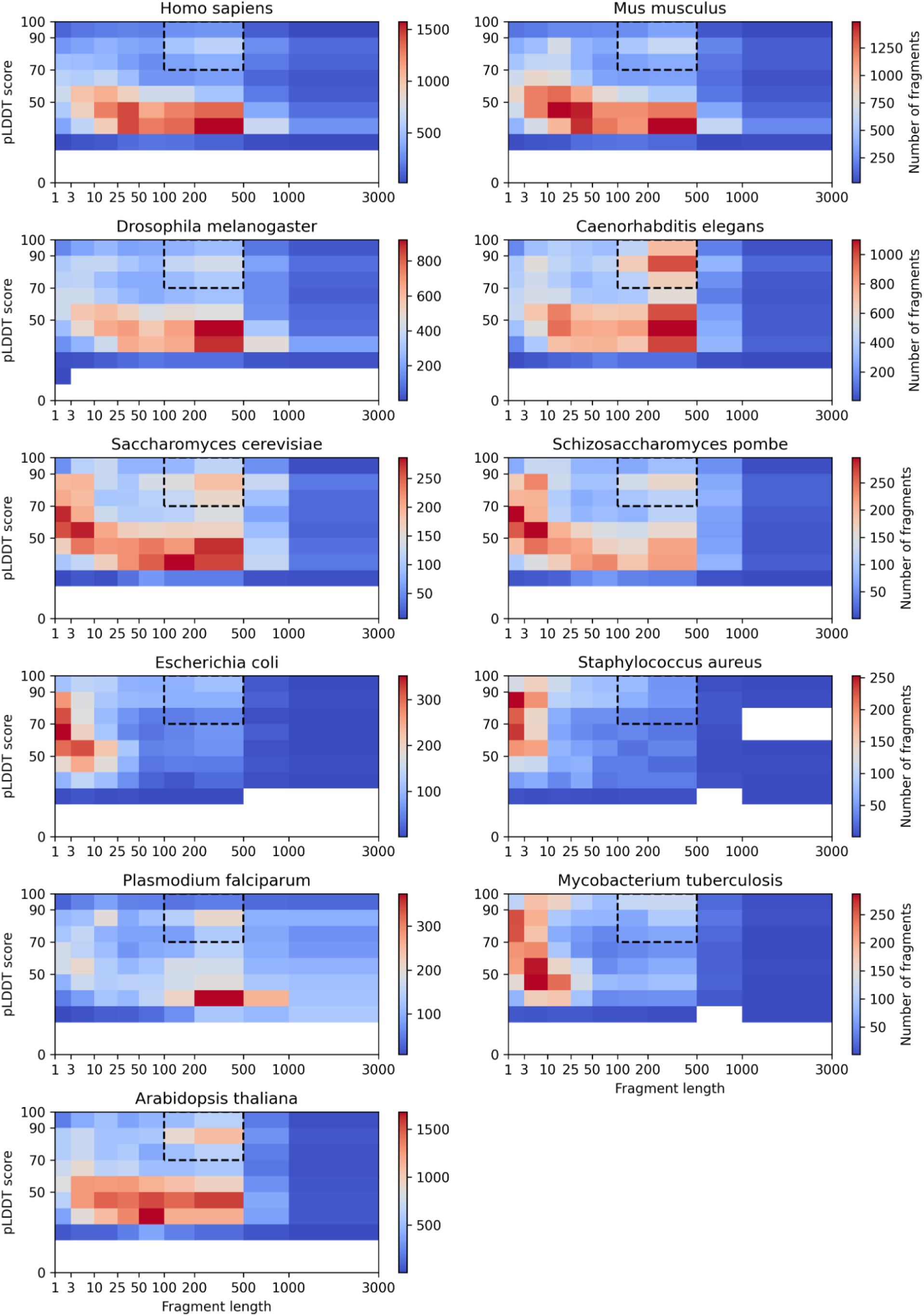
Distribution of median pLDDT scores vs. Fragment length. Only fragments less than 3000 are shown. For most proteomes, we find some level of enrichment for high confidence fragments of length between 100 to 500 amino-acids (boxed areas).

**Fig S4.**
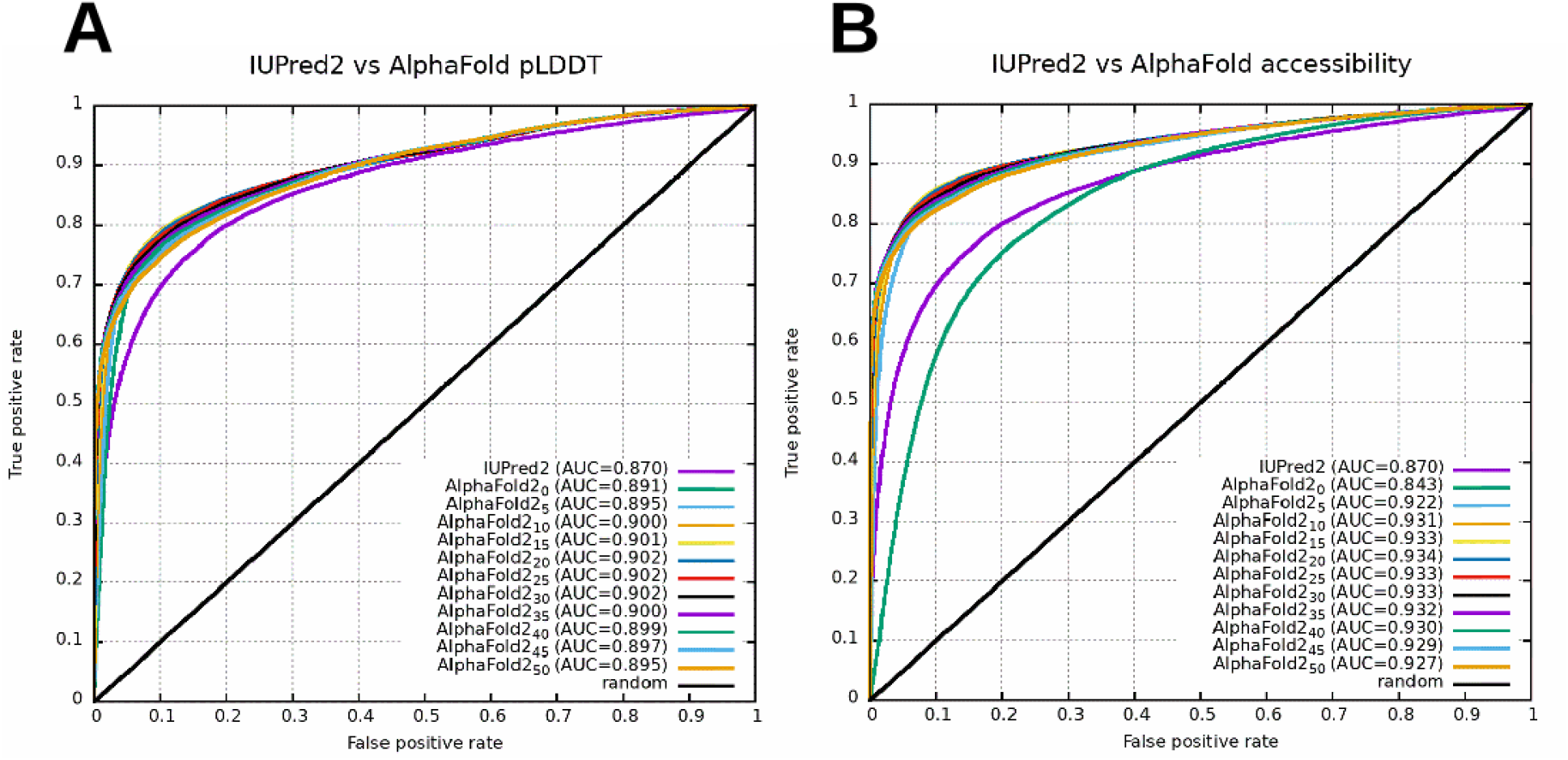
Benchmarking AF2-derived measures against IUPred2. ROC curves comparing the pLDDT confidence scores. A) and calculated relative solvent accessible surface areas. B) of AF2 structure predictions to IUPred2. Indices in legends mark the window sizes used to smooth the values along the sequence. AUC = area under the curve, the overall measure of performance, where a perfect prediction is AUC=1.0 and a random prediction is AUC=0.5.

**Fig S5.**
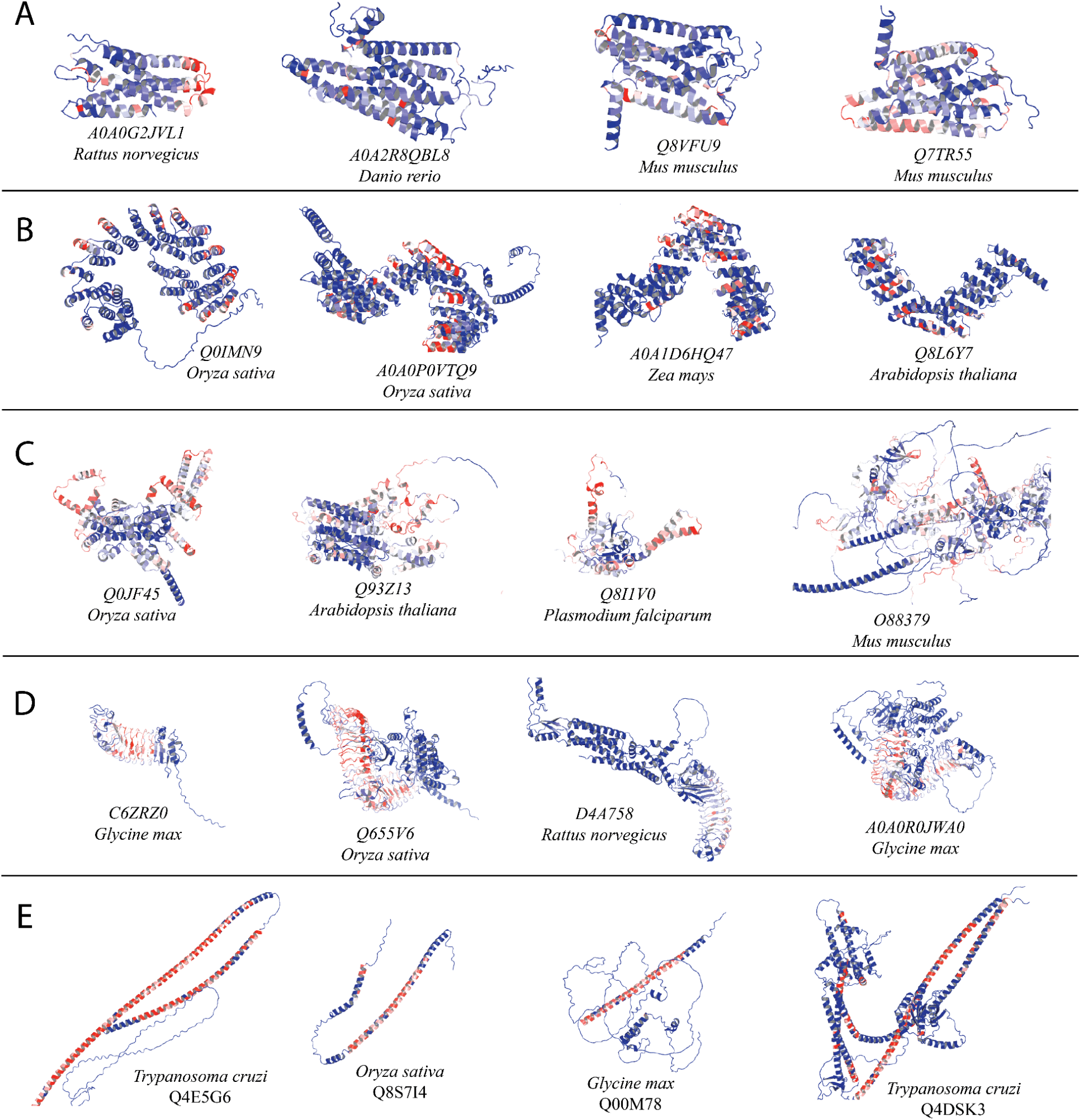
Representative structures from topics representing likely rare or new folds. Residues are colored according to their contribution to the topic under consideration - red residues have the highest contribution, while blue residues are specific to the example and not to the topic. A) GPCR olfactory receptors. B) Plant pentatricopeptide repeat proteins. C) ATP- and ion-binding proteins. D) Proteins with Leucine rich repeats. E) Long α-helical constructs

**Fig S6.**
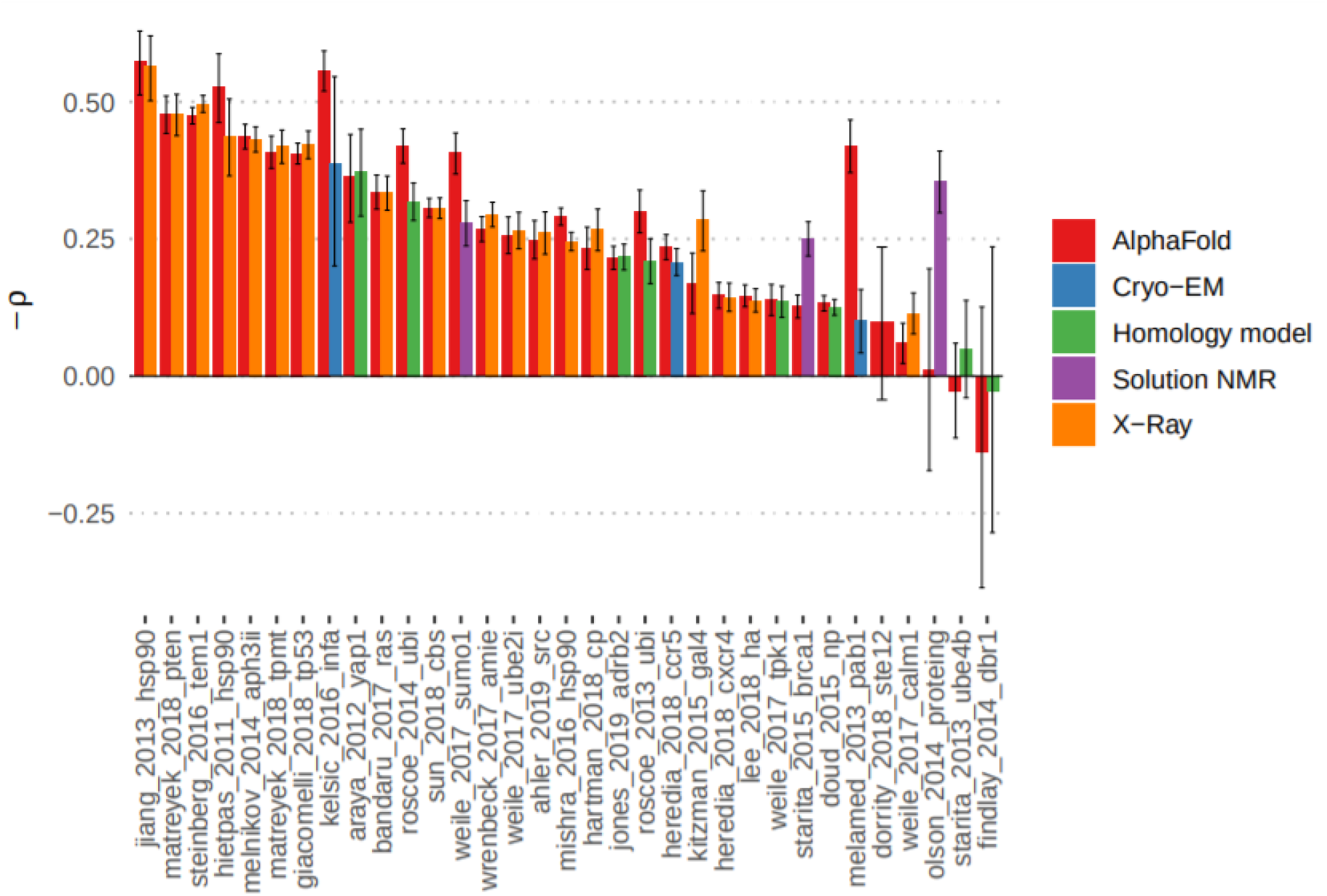
Correlation between structure based prediction of destabilisation and measured impact on protein function by deep mutational scanning experiments. Relation between the predicted ΔΔG for mutations with measured experimental impact of the mutation from deep mutational scanning data (−1*pearson correlation). The predicted change in stability was done with FoldX using structures from AF2 or available experimental models.

**Fig S7.**
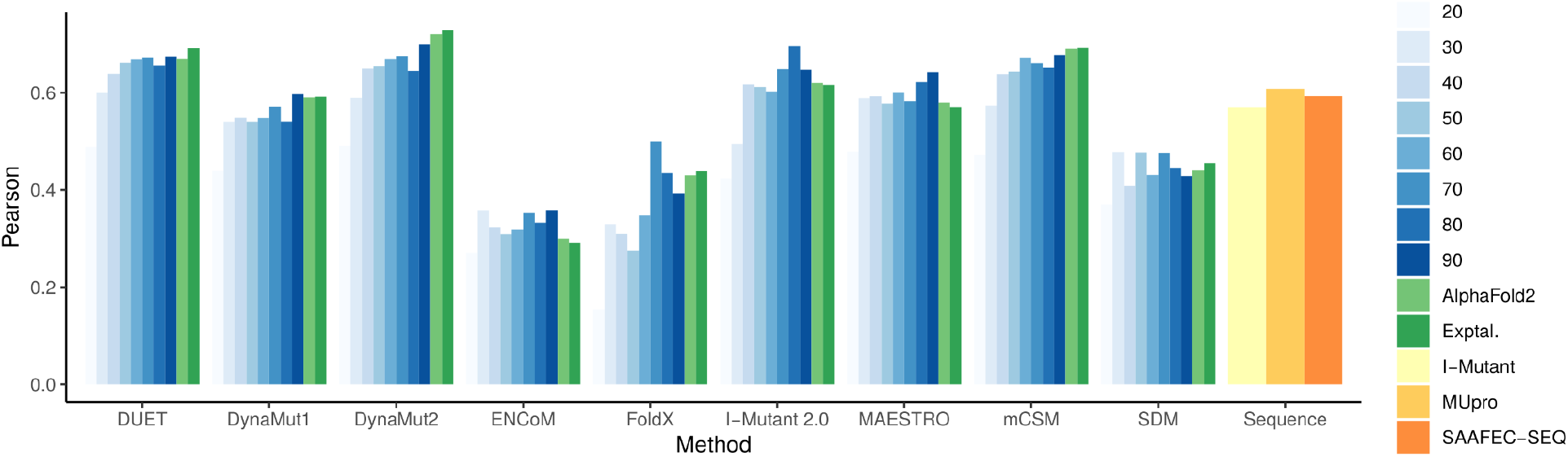
Comparative performance of methods predicting stability changes upon mutation using AlphaFold2 and MODELLER models. The performance of nine well established structure-based methods is contrasted on a mutation benchmark data set when presented to either AlphaFold2 models (light green bars) or homology models (blue bars) derived from templates of varying identity levels. Baseline performance using experimental structures (dark green bar) and of three sequence-based tools (yellow and orange bars) are also shown.

**Fig S8.**
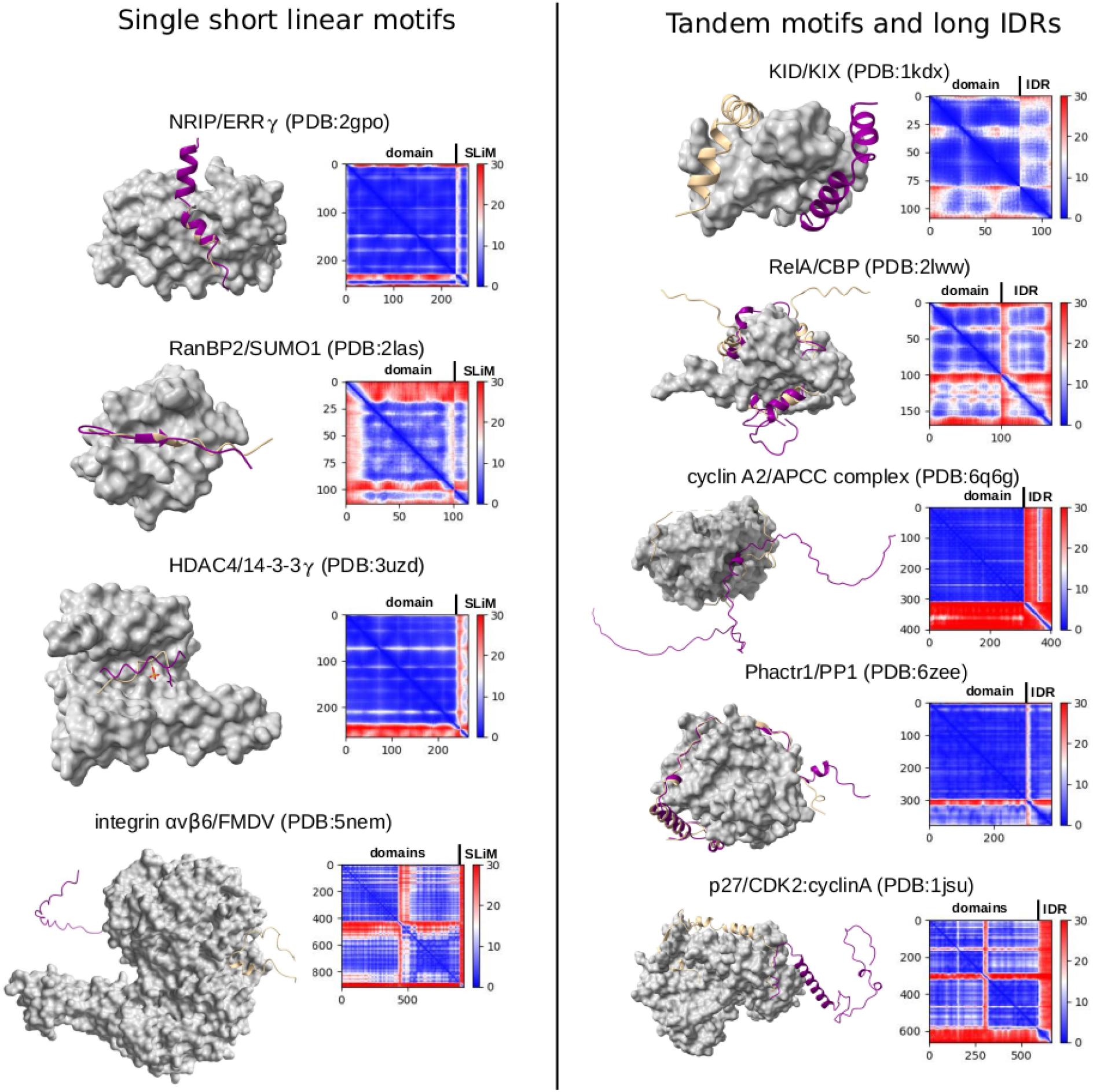
AF2-predicted complex structures of single IDRs bound to ordered partners. Left - short linear motifs (SLiMs), from top to bottom: NRBOX motif in NRIP1 bound to ERR3, RanBP2 SIM bound to SUMO, HDAC4 14-3-3 phosphomotif bound to 14-3-3gamma, and the RGDLxxL motif of the FMDV viral protein bound to integrin alphavbeta6. Right — long IDRs with several binding regions, from top to bottom: KID region of CBP bound to the KIX domain, TAD of RelA bound to the TAZ domain of CBP, Cyclin-A2 bound to Cdc20, Phactr1 bound to PP1 and p27 bound to the CDK2:cyclinA complex.

**Fig S9.**
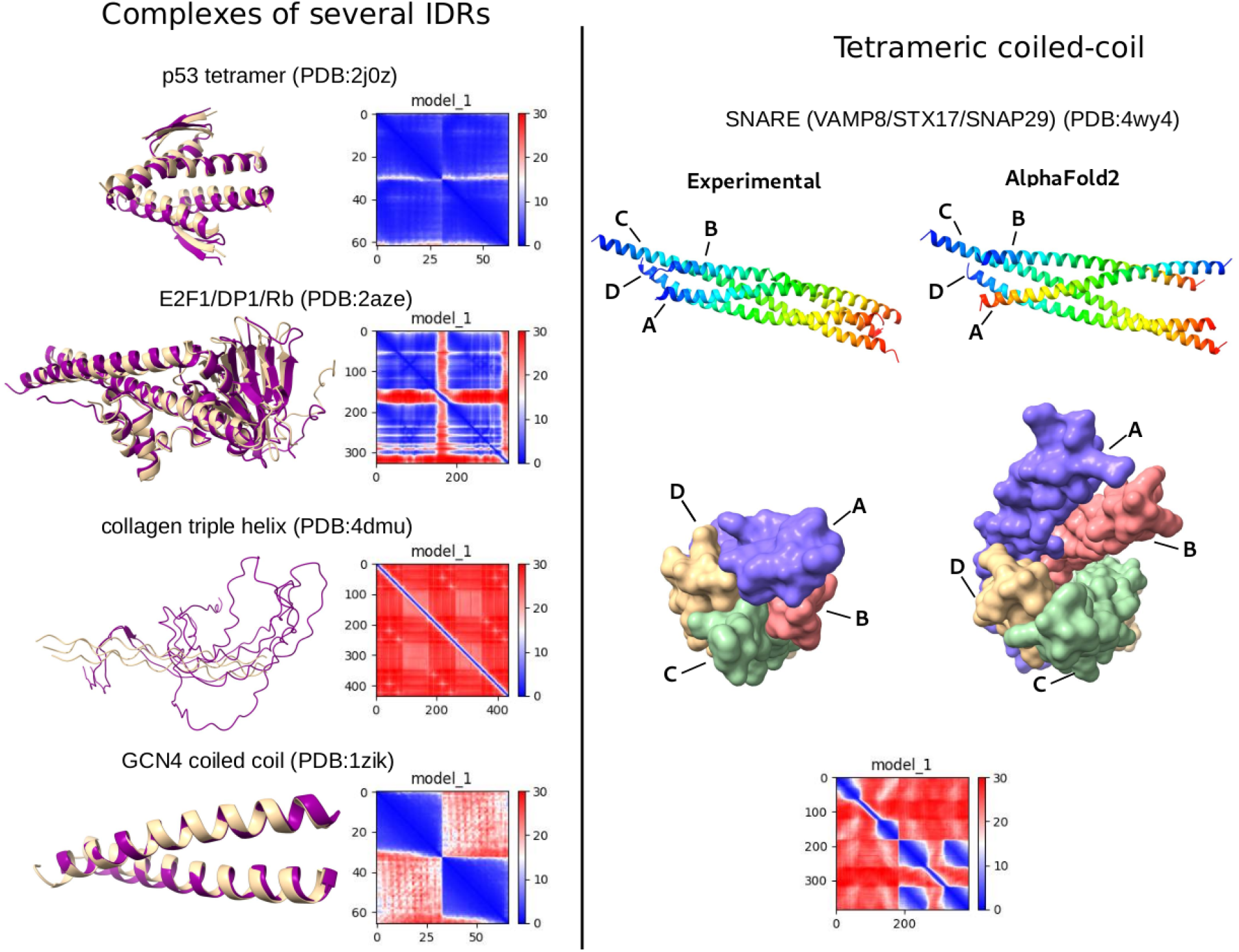
AF2-predicted complex structures of complexes with multiple IDRs. Left — IDR-only complexes, from top to bottom: p53 tetramerization region, Rb:E2F1:DP1 heterotrimer, collagen triple helix and the GCN4 prototypic leucine zipper. Right: Autophagic SNARE core complex (Vamp8 / Syntaxin-17 / SNAP29).

## Author Contributions

EPP, AV, VRS performed the AlphaFold and RoseTTaFold comparison. AOZ performed the AlphaFold and SwissModel Repository comparison. JD and MA did the characterisation of structural elements in AlphaFold DB (shape-mer analysis). NED and BM performed the disorder region and disordered complex structure prediction. TIC performed the experimental model building studies. SO performed the homo-oligomeric state predictions. JJ and PeB did the pocket prediction analysis. AE, AdS, GP, PaB, WZ and PK performed the protein-protein interaction modelling. JMT, NB and RAL performed the template search study. ASD, PeB, AmS, KLL, LLG, CHMR, DBA, DEVP performed the variant effect prediction analyses. DB provided technical assistance with running AF2 predictions. AF and SV contributed with analysis suggestions and text revision. PeB, AV, AmS, KLL, NED, TIC, SO, AE, JMT, JD, DBA supervised work. All authors contributed to the writing of the manuscript.

## Acknowledgements

B.M. has received funding from the European Union’s Horizon 2020 research and innovation programme under the Marie Skłodowska-Curie grant agreement no. 842490 (MIMIC). AmS received funding from the Lundbeck Foundation (R272-2017-4528) and Novo Nordisk Foundation (NNF18OC0033950). DBA received funding from the National Health and Medical Research Council of Australia (GNT1174405) and the Victorian Government’s Operational Infrastructure Support Program. KLL received funding from the Novo Nordisk Foundation (NNF18OC0033950). DEVP received funding from an Oracle Research Grant. AOZ received funding from the Russian Science Foundation (RSF) 20-14-00121. AE, AdS, GP, PaB, WZ received funding from the Swedish Research Council for Natural Science, grant No. VR-2016-06301 and Swedish E-science Research Center. TIC received funding from the Wellcome Trust.

